# Drude SILCS-Nucleic: Harnessing Explicit Electronic Polarization in Targeting RNA and DNA for Drug Design

**DOI:** 10.64898/2025.12.20.695737

**Authors:** Haley M. Michel, Anne M. Brown, Alexander D. MacKerell, Justin A. Lemkul

## Abstract

The growing interest in nucleic acids as therapeutic targets has prompted the development of novel computational methods to facilitate drug discovery. In this study, we extend the Site Identification by Ligand Competitive Saturation (SILCS) methodology to characterize ligand–nucleic acid interactions using the Drude polarizable force field. We demonstrate the ability of the Drude force field to better model solute-nucleic acid interactions, resulting in improved identification of known binding sites and ligand binding favorability predictions across a diverse set of nucleic acid structures. This new workflow addresses limitations in previous SILCS studies by exploiting the enhanced sampling of solutes in the original SILCS-RNA workflow, accurately modeling the interactions of charged species, and improving solute sampling in minor groove binding sites. These results establish this Drude-based SILCS workflow as a valuable tool for structure-based drug design targeting nucleic acids and offer insights into solute preferences that can guide rational ligand design.

## Introduction

Nucleic acids are central to the structure, function, and regulation of the human genome and can adopt complex three-dimensional structures with distinct functions. Our understanding of these functional roles continues to expand, highlighting the involvement of DNA and RNA structures in the regulation of biological processes. Many DNA and RNA structures, such as riboswitches and G-quadruplexes, have been identified as key regulators of gene expression, connecting them to the progression of bacterial infections, cancer, and human neurodegenerative diseases.^1–4^ Consequently, nucleic acid structures have emerged as a promising therapeutic avenue, offering an alternative to traditional protein-based drug design, which is often limited by mutations and off-target effects.

Over the past 30+ years, there has been a concerted effort to develop and integrate computational methods into the drug discovery process; however, the majority of such methods have been created and optimized for proteins. Although recent advances in protein–ligand interaction prediction have shown considerable success in protein targeting, extending these methods to nucleic acids remains challenging due to their unique structures and highly charged electrostatic nature. To address this gap, several nucleic acid specific molecular docking methods such as HADDOCK^5^ and RLDOCK,^6^ have been developed. Nevertheless, their success rates remain low, largely because they rely on static structures and insufficiently account for electrostatic effects.^7^

To incorporate structural dynamics, so-called “co-solute” methods have emerged, em-ploying molecular dynamics (MD) simulations to identify favorable interaction sites of small probe molecules, co-solutes, on the target structure. Currently, only two such methods exist for RNA: SILCS-RNA^8^ and the more recent SHAMAN,^9^ each using distinct sets of co-solutes to probe potential binding sites on nucleic acids. The Site Identification by Ligand Competitive Saturation (SILCS) method^10–12^ combines Grand-Canonical Monte Carlo and MD (GCMC-MD)^13^ to enhance the sampling of water and solutes as they compete for interaction sites on the target. The resulting interaction patterns are assembled into “FragMaps,” which are converted to grid free energy (GFE) values representing the affinity of functional groups throughout the target molecule. FragMaps can be used to identify binding “hotspots,”^14^ guide ligand relaxation or docking in the field of FragMaps,^15^ estimate binding affinity,^15,16^ and generate pharmacophores^17^ for virtual screening applications.

The SILCS-RNA protocol has shown promise in predicting affinities of known ligands to their corresponding binding sites including information on the contribution of individual functional groups to binding. However, the use of the nonpolarizable CHARMM force field (FF)^18,19^ in the standard SILCS protocol limits the accurate modeling of interactions involving charged species. As a nonpolarizable model, the CHARMM FF cannot account for changes in electronic structure in response to the local environment, resulting in a limitation that is an important contribution for highly charged targets such as nucleic acids.

Polarizable FFs, such the Drude polarizable FF^20^ and the AMOEBA FF,^21^ provide a viable solution to this challenge.^22^ The Drude FF explicitly incorporates the polarization response in the functional form of the FF and is based on the classical Drude oscillator model. Within Drude, negatively charged particles are attached to all non-hydrogen atoms by a harmonic spring, allowing the atomic dipoles to dynamically respond to the surrounding electric field. This polarization response is critical for capturing the nuanced interactions that drive RNA-ligand binding, including hydrogen bonding, *π*-stacking, and electrostatic complementarity.

The Drude FF has previously been applied alongside the SILCS method to evaluate the role of electronic polarization in protein-solute interactions.^8^ The use of Drude improved the modeling of solute-target interactions, as variations in induced dipole moments led to better-defined interaction orientations and enhanced solute occupancy and favorability within known binding sites. However, this study was limited to proteins and was conducted without GCMC-MD, potentially reducing solute sampling compared to the standard non-polarizable SILCS workflow. Extending such studies to nucleic acids is essential, as their uniform and large charge density poses fundamentally different electrostatic challenges than those encountered in proteins.

In this study, we extend the original SILCS-RNA workflow to incorporate the Drude polarizable FF and assess its performance across a diverse set of DNA and RNA structures. This approach leverages electronic polarization and the enhanced solute sampling of the nonpolarizable simulations to more accurately capture solute-nucleic acid interactions. We evaluate its ability to model the complex electrostatic environment of nucleic acids and to predict the binding sites and poses of known ligands. Furthermore, we compare the performance of this method with established docking programs to identify current limitations in nucleic acid-ligand interaction prediction. Overall, this work represents a critical step toward improving the accuracy of computational approaches for nucleic acid-targeted drug discovery.

## Methods

The Drude SILCS-Nucleic workflow was applied to six nucleic acid systems for which small molecule-RNA interaction data are available. All systems are listed in Table 1, along with the corresponding Protein Data Bank (PDB) accession codes and relevant details such as associated ions and ligands. The selected systems form a robust test set encompassing both DNA and RNA, a range of tertiary structures, essential ions, and both single- and double-stranded forms (Table 1 and Figure S1). The four RNA systems were previously investigated using the SILCS-RNA protocol^8^ and the CHARMM36 (C36) FF,^18,19^ enabling direct comparison and assessment of reproducibility between the original nonpolarizable results and the updated, polarizable workflow.

**Table 1:**
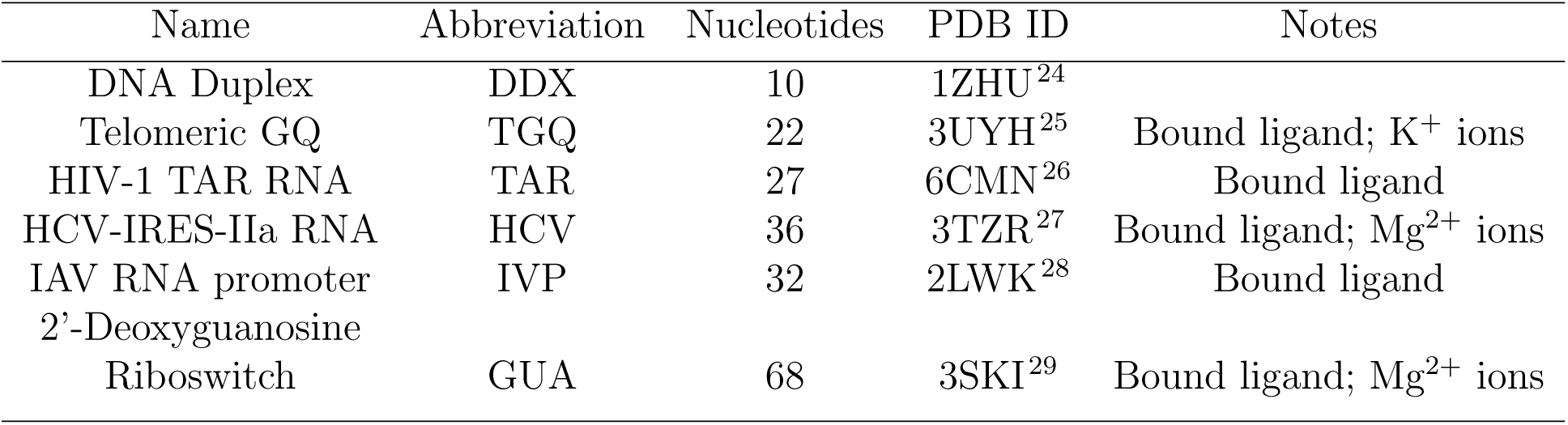
Drude SILCS-Nucleic system information.

| Name | Abbreviation | Nucleotides | PDB ID | Notes |
| --- | --- | --- | --- | --- |
| DNA Duplex | DDX | 10 | 1ZHU<sup>24</sup> |  |
| Telomeric GQ | TGQ | 22 | 3UYH<sup>25</sup> | Bound ligand; K<sup>+</sup> ions |
| HIV-1 TAR RNA | TAR | 27 | 6CMN<sup>26</sup> | Bound ligand |
| HCV-IRES-IIa RNA | HCV | 36 | 3TZR<sup>27</sup> | Bound ligand; Mg<sup>2+</sup> ions |
| IAV RNA promoter | IVP | 32 | 2LWK<sup>28</sup> | Bound ligand |
| 2'-Deoxyguanosine<br>Riboswitch | GUA | 68 | 3SKI<sup>29</sup> | Bound ligand; Mg<sup>2+</sup> ions |

The Drude SILCS-Nucleic workflow is an extension of the original C36 SILCS-RNA protocol, in which the systems are subjected to 100 cycles of 200,000 steps of non-equilibrium oscillating chemical potential Grand Canonical Monte Carlo (GCMC) simulations followed by subsequent 1-ns MD simulations to enhance the sampling of solutes. The Drude SILCS-Nucleic workflow uses 100 snapshots extracted from all 10 of the C36 systems, evenly spaced in time, to benefit from the enhanced sampling of the SILCS-GCMC process with respect to the sampling of the water and solute distributions (Figure 1).

**Figure 1:**
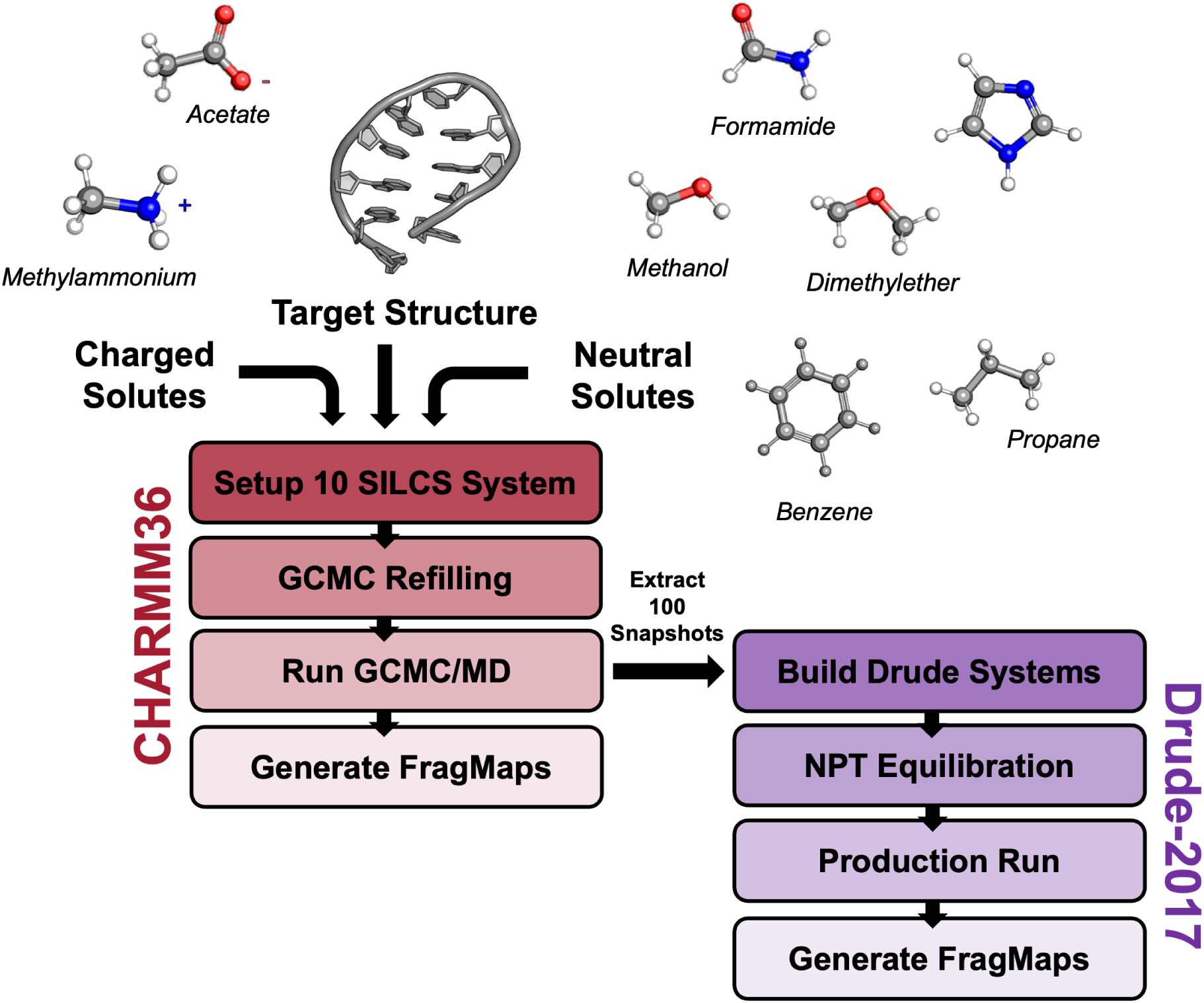
Drude SILCS-Nucleic workflow with the Drude FF.

### Nucleic Acid Structure Preparation

To start the SILCS simulations, 3D structures of each system were obtained. RNA structures (TAR, HCV, IVP, and GUA) were obtained from Kognole *et al.* to ensure the same starting structures from the development of the SILCS-RNA method.^8^ The two DNA structures (DDX and TGQ) were obtained from the PDB and prepared using the CHARMM program,^3^ following the methods of our previous work.^31–33^ The minimized DNA systems were then stripped of water and ions, except for the core K^+^ ions within TGQ, and used as the starting point for the additive SILCS-RNA simulations.

### Additive SILCS-RNA Workflow: System Setup

The SILCS-RNA protocol creates two sets of SILCS simulations, one containing neutral solutes: benzene (BENX), propane (PRPX), dimethyl ether (DMEE), methanol (MEOH), formamide (FORM), imidazole (IMIA), and one containing charged solutes: acetate (ACEY) and methylammonium (MAMY). This approach allows all neutral solutes to compete with each other and water for interaction sites without interference from the highly-favorable interactions of MAMY and the negatively charged phosphate backbone of the target structure.^8^ Once PDB files were obtained, GRO-MACS version 2022.5^34^ was used to generate the nucleic acid topology according to the C36 FF. Then, ten independent systems were created for each set by placing the nucleic acid in a water box containing randomly placed solutes. The dimensions of the box were set to be 15Å from the nucleic acid on all sides and the solutes and water were added to concentrations of 0.25 M and 55 M, respectively.

### CHARMM SILCS Workflow: GCMC Refilling

Each system was then subjected to non-equilibrium GCMC sampling. First, a GCMC-active region was defined as a subvolume of the simulation box using the maximum x-, y-, and z-dimensions of the nucleic acid within the 15 Åmargin to the edge of the full simulation box. All water and solutes were removed from the GCMC-active region and refilled in a two-stage or three-stage process for charged and neutral systems, respectively. The two-stage process was applied to the set of charged solute systems, such that ACEY and MAMY solutes were added in stage 1 followed by the addition of water in stage 2. The three-stage process was used for the set of neutral solute systems. First, BENX, PRPX, and IMIA were inserted to a concentration of 0.35 M to ensure these larger solutes could access any buried binding pockets or intercalation sites. FORM, MEOH, and DMEE were then added to 0.25 M in stage 2 and finally water was inserted to 55 M in stage 3. Each stage of refilling involved 200,000 steps of GCMC, during which solutes were randomly inserted, deleted, translated, or rotated within the GCMC-active region using the excess chemical potential, *µ_ex_*, of each solute to accept or reject each move according to the Metropolis criterion.^13^

### CHARMM SILCS Workflow: GCMC-MD

Following GCMC refilling, each system underwent steepest descent minimization and restrained equilibration for 100 ps. The production stage involved 100 cycles of SILCS GCMC-MD consisting of four stages. First, the system underwent 200,000 steps of GCMC of all the solutes and water molecules to enhance solute sampling. The system then underwent steepest descent minimization for a maximum of 5,000 steps or upon reaching a force tolerance of 10 kJ/(mol·nm), 100 ps of MD equilibration, and 1 ns of production MD simulation. During production, weak harmonic restraints with a force constant of 50.208 kJ/(mol·nm^2^) [equivalent to 0.12 kcal/(mol·Å^2^)] were applied to the C1’ sugar atoms and the N1 atom of purines and N3 atom of pyrimidines unless otherwise noted.

For TGQ, the weak SILCS restraints were not able to retain the K^+^ ions within the GQ core. Such repulsion has been shown to be an artifact of some nonpolarizable FFs^31,32,35^ and therefore must be accounted for in these simulations. The interaction energy between the core K^+^ ions with the C36 FF was calculated as 7.3 kcal/mol (30.5 kJ/mol) in a previous study.^31^ In Drude FF simulations, K^+^ ions are retained in the GQ tetrad cores, with fluctuations of ±0.3 Å (0.03 nm). Thus, the strength of the restraint must be set to offset this repulsive energy and allow for some small variations in position along the *z*-axis. Assuming a harmonic oscillation of these ions such that 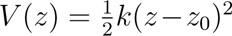, a force constant of 67874 kJ/(mol·nm^2^) [154.86 kcal/(mol·Å^2^)] will produce this behavior. As such, this value was used as the force constant for position restraints applied to K^+^ within the TGQ core. A total simulation time of 1 *µ*s was thus obtained across the 10 independent systems for both neutral and charged systems. All GCMC-MD simulations were conducted using SilcsBio version 2022.2 (SilcsBio LLC) in conjunction with GROMACS version 2022.5.^34^

### CHARMM SILCS Workflow: FragMap Generation

Once the SILCS simulations were completed, selected solute atoms are categorized based on their functional group class (Supplementary Figure S1). The selected solute atoms were then binned into 1-Å^3^ cubic voxels extracted from all the trajectories from the GCMC-MD simulations to calculate the respective probability distributions. The distributions were then normalized with respect to the number of frames from the MD simulations, total number of atoms of the selected atom type in each solute (n_atoms_)(eg. 6 aromatic carbons in benzene), and the bulk occupancy (OCC_bulk_), which is defined by the average number of each solute relative to the number of waters with water being assigned a concentration of 55 M. These values were converted to free energies, termed grid free energies (GFE) using a Boltzmann transformation as shown in Eqs. 1 and 2:

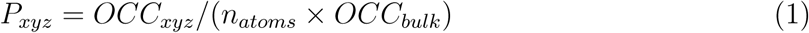

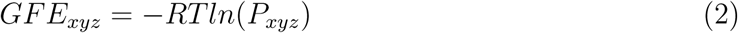

The resulting grids are referred to as the GFE FragMaps. FragMap convergence was evaluated via an overlap coefficient (OC), which determines the extent of solute sampling overlap between the first and last sets of five independent simulations. That is, the OC quantifies how similar the probability distributions are from simulations 1-5 and 6-10, according to Eq. 3:

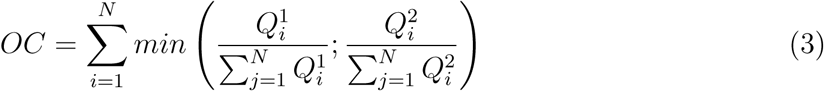

where *N* is the number of voxels in the FragMaps and 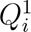 and 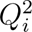 are the occupancies of voxel *i* for set 1 (simulations 1-5) and set 2 (simulations 6-10). OC values range between 0 and 1, with 0 signifying completely different FragMaps and 1 signifying equivalent FragMaps. OC values ¿ 0.6 have been shown to be adequately converged.^13^ Exclusion maps were also obtained to define the volume excluded from the solute and water non-hydrogen atoms by the presence of the nucleic acid. FragMaps were visualized using the SilcsBio FragMap Plugin with PyMOL.^36^

### Drude SILCS-Nucleic Workflow

Prior to development of the Drude SILCS-Nucleic Workflow, the Drude topology for formamide was evaluated and refined based on experimental data (Supplementary Methods S1, Table S2, and Listing S1). To obtain the benefits of the GCMC sampling conducted during the CHARMM simulations, ten snapshots from each of the ten independent C36 systems were extracted from both the charged and neutral solute sets. These snapshots were taken from the end of cycles 10, 20, 30, 40, 50, 60, 70, 80, 90, and 100 of the C36 systems. Thus, each of the ten nonpolarizable systems contributed ten initial states to the Drude simulations, resulting in 100 systems for each set. As with the C36 systems, the neutral and charged solutes were treated in separate simulation sets for the Drude systems.

### Drude SILCS-Nucleic Workflow: System Construction

To build the Drude systems, the coordinates from the last frame of each selected C36 cycle were saved in PDB format. These PDB files were converted to CHARMM-compatible coordinate files using an in-house Python script to add segment identifiers and adjust atom naming to be compliant with the Drude FF and the CHARMM program. Negatively charged Drude oscillators were added to all non-hydrogen atoms in the system and lone pairs were added to all hydrogen bond acceptors. TIP3P^37^ water molecules were converted to the polarizable SWM4-NDP^38^ model, and the Drude-2017 FF^39–41^ was applied to generate a new Drude topology. The Drude systems then underwent a 1000-step steepest-descent minimization followed by 2000 steps of Adopted-Basis Newton-Raphson minimization with all real atoms restrained to relax the Drude oscillators, therefore allowing the induced dipoles to relax in the field of the fixed charges.

### Drude SILCS-Nucleic Workflow: Equilibration

Equilibration was performed for 1 ns under an *NPT* ensemble in OpenMM. Position restraints were applied to all non-hydrogen atoms of the nucleic acids and any bound cations in the system with a force constant of 500.0 kJ/(mol·nm^2^) [1.195 kcal/(mol·Å^2^)]. The solute and water atoms were not restrained during equilibration, allowing them to relax around the nucleic acids. The temperature was set to 298 K and maintained with the Langevin thermostat method,^42^ using a friction coefficient of 5 ps^-1^ while the Drude oscillators were coupled to a low-temperature thermostat at 1 K with a friction coefficient of 20 *ps^−^*^1^. The pressure was maintained at 1 atm throughout equilibration by a Monte Carlo barostat. The van der Waals potential was switched from 10 to 12 Å and the particle mesh Ewald method^43,44^ was used to calculate electrostatic interactions with a real-space cutoff of 12 Å and 1-Å Fourier grid spacing. To avoid polarization catastrophe, a “hard wall” constraint^45^ was applied to maintain the Drude-atom bond lengths within 0.2Å.

### Drude SILCS-Nucleic Workflow: Production Simulations

Equilibrated systems were then used for production simulations. To match the nonpolarizable simulations, weak po-sition restraints were applied to the C1’ sugar atoms and the N1 atom of purines and N3 atom of pyrimidines with a force constant of 50.208 kJ/(mol·nm^2^) [0.12 kcal/mol·Å^2^)]. These restraints maintained the global conformation of the nucleic acids while allowing local structural fluctuations. In contrast to the nonpolarizable simulations, no restraints were applied to the K^+^ ions in the TGQ system, given that the Drude FF more accurately models the coordination of these ions.^31,32,35^ For each system cycle, 20 ns of production MD was conducted in OpenMM resulting in a total of 200 ns of simulation time for each of the ten starting conformations in each solute set. Thus, a total of 2 *µs* of sampling was obtained for each nucleic acid system.

### Drude SILCS-Nucleic Workflow: FragMap Generation

FragMaps for the Drude simulations were generated using a similar process as with C36. The SilcsBio scripts required updates to read in Drude file naming and Drude FF atom types. For each of the 100 systems in both solute sets, the 20-ns trajectory was oriented using LOOS^46,47^ to ensure proper periodic imaging and that the nucleic acid structure was centered at the Cartesian origin. A similarly oriented PDB file was also generated for each of the 200 systems using LOOS. Occupancy maps were produced for every solute across all production simulations. These maps were then collated per solute to generate a single map for each solute, representing all sampling across the entire 2 *µs* simulation time. GFE values were generated for each map and the OC was calculated. Due to the approximate nature of normalizing the GFE values, the Drude FragMaps required an offset of ∼1.2 - 1.3 kcal/mol to reach a bulk GFE = 0 kcal/mol. This offset was applied using an in-house script for visualization in PyMOL.

### SILCS-MC and LGFE Calculations

The SILCS-MC program was used to exhuastively dock or locally refine (pose-refinement) ligand poses and determine the Ligand Grid Free Energies (LGFE, corresponding to the predicted binding affinity) from the atomistic GFE contributions to ligand binding for the TGQ, TAR, HCV, IVP, and GUA system. The crystal or NMR structure of each ligand-bound complex, if not already being used for our SILCS simulations, was aligned with the structures used for our SILCS simulations to approximate the binding site. Ligands were prepared in Schrödinger Maestro where Epik^48^ was used to calculate pK_a_ values and protonation states. SILCS-MC uses the CHARMM General Force Field (CGenFF)^49^ to assign atom types that are then categorized with respect to FragMap types for selected atoms of the ligands within the SILCS atom classification scheme.^15^ This classification was then used to calculate the LGFE of the ligands based on the overlap of the classified atoms with the corresponding FragMaps. The LGFE plus the CGenFF intramolecular energy were used for the Metropolis criteria using MC sampling of the ligand conformation and orientaton to obtain the most favorable ligand conformation with respect to FragMaps and Exclusion map. Two distinct SILCS-MC protocols were used; for local pose refinement and exhaustive docking with minor adjustments to parameters as needed. The protocols used for each ligand can be found in Supplemental Table S5.

### SILCS-MC Local Pose Refinement

The default pose-refinement protocol for SILCS-MC consisted of up to 50 rounds of Monte-Carlo sampling followed by Simulated Annealing (MC/SA) within a sampling sphere with a 1-Å radius from the center-of-geometry of the input ligand pose. Each cycle of MC/SA consisted of 100 steps of MC at 300 K, to randomly sample diverse conformations and orientations, followed by 1000 steps of SA from 300 to O K to identify the most favorable local pose. Each cycle began at the pose of the input ligand and MC sampling was limited to translations with a maximum step size of 0.5 Å, molecular rotations with a maximum size of 15°, and intramolecular dihedral rotations with a maximum step size of 180°. The final pose from the MC stage was then used as the starting point for the SA sampling. SA sampling employed tighter configurational restraints, only allowing a maximum translation step of 0.2 Å, a maximum molecular rotation of 9°, and a maximum dihedral rotation of 9°. Convergence of the MC/SA simulations was reached when the LGFE score among the top three ligand poses over the indivudal rounds of pose refinement was within 0.5 kcal/mol of one another up to a maximum of 50 rounds. The three lowest-LGFE poses were then selected and written to a multi-SDF file. Final scores are based on only the LGFE values.

### SILCS-MC Exhaustive Docking

For the SILCS-MC Docking protocol, five independent jobs were generated for each ligand. The general methodology for docking was the same as for pose refinement except exhuastive docking performed up to 250 rounds of MC/SA, with 10,000 steps of MC at 300 K and 40,000 steps of SA from 300 to 0 K. Each round started with a reoriented ligand pose within the sampling sphere. The docking protocol employed more permissive MC moves, allowing maximum translations up to 1 Å, maximum molecular rotations of 180°, and maximum intramolecular dihedral rotations of 180°. For SA, the same maximum allowances were used as in pose refinement. The lowest-LGFE poses were again written to a multi-SDF file, then separated into individual SDF files, and used for visualization. Root-mean-squared deviation (RMSD) was calculated for the ligand poses using the rms tool from OpenBabel.^50^ The SDF files from both the experimental and the SILCS-MC refined poses were used as the inputs for this calculation.

Schrödinger Jaguar was used to predict the pK_a_ values of the nitrogen atoms in the *N* - methylpiperazine and morphilino rings. The pK_a_ values for these two groups were determined to be 8.07 and 8.41, respectively, indicating that both nitrogen atoms are predominantly protonated at pH 7 and were therefore treated as such.

### Molecular Docking with Glide and Gnina

Schrödinger Maestro^51^ was used to perform docking of the C36 and Drude structures of all nucleic acid systems. From the PDB files, “POT” (K^+^ ions in C36 and Drude nomenclature) were incorrectly interpreted by Maestro as phosphorus (P) and needed to be edited manually to ensure correct charge assignment. Structures and ligands were prepared using Protein Prep and LigPrep to add hydrogen atoms and ensure the appropriate protonation state of each ligand and obtain .mae format files for docking. The receptor grids were generated around each binding site with inner box dimensions set to encompass all core ligand atoms and the outer box dimensions set to contain all ligand atoms. The generated receptor grid contained information about the shape and interaction properties of the receptor. The ligand was treated as flexible during the docking process to generate new conformations based on rotatable bonds. A series of filters were used to search for “site points” within the binding site, which represent potential ligand binding sites. The sites were evaluated based on how close the ligand was to the receptor and how well the ligand orientation fit in the site. The resulting ligand interactions with the receptor were scored using the ChemScore empirical scoring function^52^ that evaluates favorable hydrophobic, hydrogen-bonding, and metal-ligation interactions while penalizing steric clashes. Top scoring poses were then re-scored after being refined, during which the ligand was allowed to move by ±1 Å in x-, y-, or z-directions. The best refined poses underwent energy minimization using the OPLS4 FF^53^ and the van der Waals and electrostatic information within the receptor grid. The minimized poses were then re-scored using the GlideScore function and a maximum of 9 poses were obtained.

Gnina version 1.3^54,55^ was also used to dock the ligands with box sizes of at least 20 Å × 20 Å × 20 Å. Since Glide uses a two box approach, the box sizes for Gnina are not exactly the same as Glide. However, the known ligand binding site served as a reference for each sampling box across all methods resulting in the same x-, y-, and z-centers being utilized allowing for each docking program to sample a similarly sized region. Within the box, the ligand underwent MC sampling to randomly rotate or translate the ligand or adjust the ligand torsional angles. After MC, the ligand conformation was minimized using a precalculated grid containing each atom type within the ligand and subsequently scored using the convolutional neural network in Gnina, after which the top nine poses were obtained.

All poses from Glide, Gnina, and SILCS-MC were visualized using PyMOL3. OpenBabel was used to calculate the RMSD between the heavy atoms of each generated pose and the experimental pose to compare each method. The dimensions of each ligand sampling region used for all methods can be found in Supplementary Tables S6-S7.

## Results and Discussion

Despite the growing recognition of nucleic acids as therapeutic targets, computational methods for modeling their interactions with small molecules remain underdeveloped compared to those for proteins. The Drude SILCS-Nucleic workflow presented here addresses this gap by explicitly accounting for polarization effects that are critical in the charged environment of nucleic acids. In the following sections, we evaluate its performance across a diverse set of RNA and DNA systems and discuss its implications for improving computer-aided drug design targeting nucleic acids.

### Impact of Electronic Polarization on Solute–Nucleic Acid Interactions

Improvements in the representation of electrostatic interactions with the Drude polarizable FF were expected to enhance interaction sensitivity and solute distributions across the nucleic acid structures, consistent with previous observations with protein targets.^8^ To assess this in nucleic acids, FragMaps were generated for both C36 and Drude systems, and simulation convergence was evaluated using overlap coefficients (OC) to quantify solute sampling consistency. OC values ≥0.6 or higher indicate adequate convergence, criteria that was met for all systems with both FFs. The Drude simulations, however, consistently yielded higher OC values than C36, by an average of 0.03 (Supplementary Tables S3 and S4). This slight increase in OC suggests that inclusion of electronic polarization resulted in refined solute placement from the inherited C36-derived starting structures, although congtributions from changes on the simulation protocol cannot be excluded.

Having established adequate convergence for both C36 and Drude, FragMaps for each nucleic acid system were visualized and compared to determine solute sampling and pattern of favorable interactions. To examine FF differences, the DNA hairpin (DDX) was analyzed as a baseline due to its simple secondary structure (Figure 2). Compared to C36, Drude simulations showed favorable FragMaps more concentrated in the centers of both the major and minor grooves, and increased enrichment of hydrogen bond donors and acceptors. These effects may be attributed to the explicit inclusion of electronic polarization, allowing both solute and target nucleic acid dipole moments to adapt dynamically to their environment allowing for more favorable interactions.

**Figure 2:**
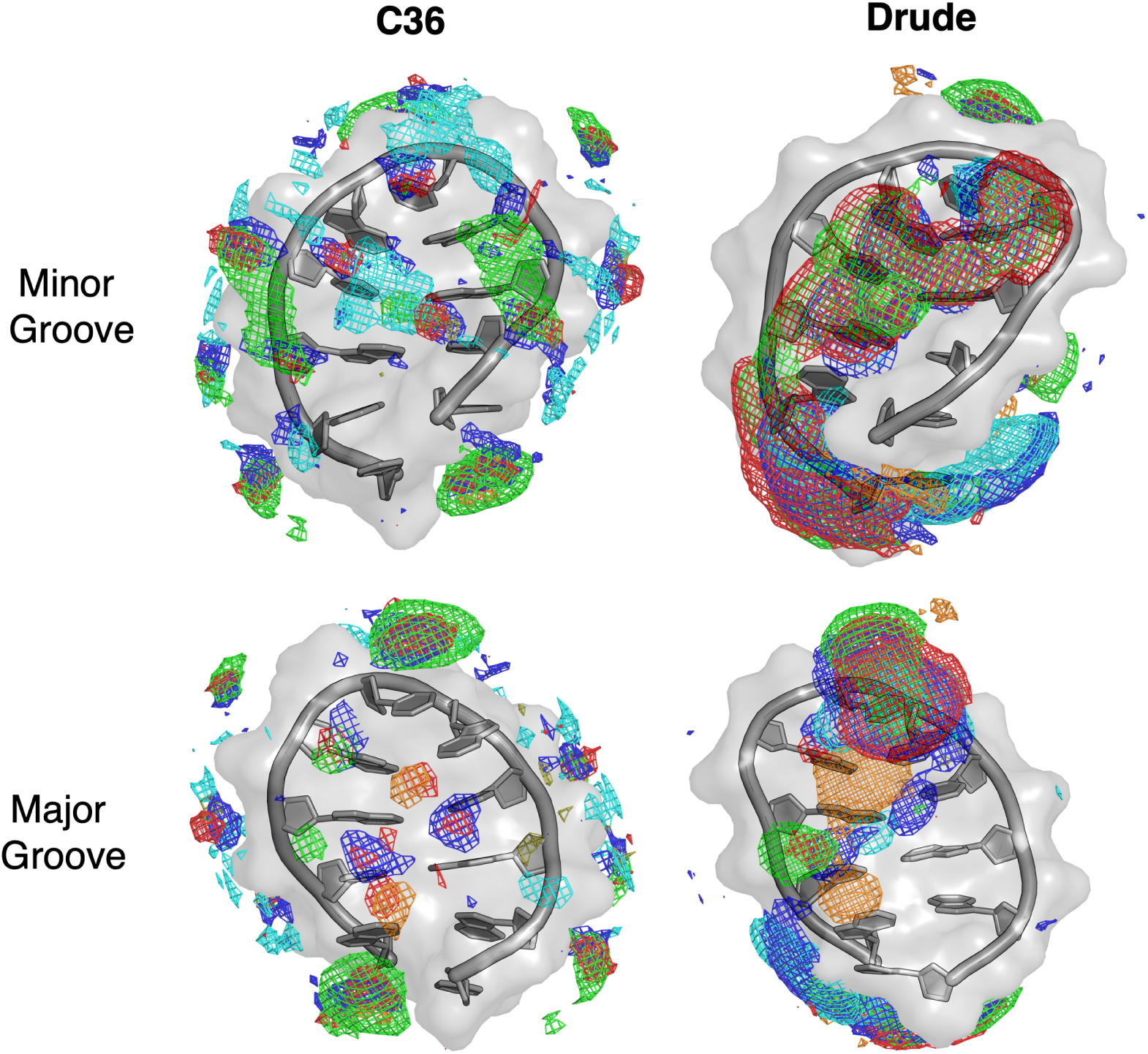
SILCS FragMaps for the DNA hairpin (DDX). All FragMaps are rendered at −0.5 kcal/mol GFE unless stated otherwise. FragMaps are shown as apolar (light green), hydrogen bond donor (blue), hydrogen bond acceptor (red), positively charged (cyan, GFE level −1.0 kcal/mol) and negatively charged (orange), and MEOO (tan).

The Drude FF also produced larger, favorable FragMap distributions than C36, particularly for acetate (orange) and apolar (green) solutes within the groove. That is, although the same chemical features were observed with the C36 FF, visualizing the larger volumes with the Drude FF at an identical GFE threshold corresponds to a broader distribution of negative GFE values, indicating a wider range of favorable solute interactions at those sites. This effect is apparent upon increasing the GFE threshold of all maps, resulting in little to no favorable FragMaps with the C36 FF and very targeted, strong favorable FragMaps with the Drude FF (Supplementary Figure S2).

The trend of larger favorable FragMap distribution and more negative GFE values with the Drude FF was consistent across all systems studied (Supplementary Figures S3 - S7), highlighting intrinsic differences between the FFs in how they model solute-nucleic acid interactions. This behavior mirrors previous findings for proteins,^8^ suggesting a general effect of polarization on solute sampling. In particular, solutes that serve as hydrogen bond donors and acceptors exhibited more pronounced sampling at favorable sites in the Drude simulations, indicating that this property is shared across proteins, DNA, and RNA. Beyond the contribution of explicit electronic polarization this outcome is likely to have contributions from the inclusion of lone pairs on all hydrogen bond acceptors, a topological feature that better accounts for the strongly directional and anisotropic nature hydrogen bonding interactions.

Given the observed difference in hydrogen bond donor and acceptor sampling between the C36 and Drude FFs, it was also important to examine how each handles interactions involving charged species. Polarizable FFs have demonstrated an enhanced ability to model charge-charge interactions compared to nonpolarizable FFs.^21,56,57^ In the previous SILCS-RNA study, strong and extensive sampling of positive charge was reported across the negatively charged RNA surfaces, driven by charge complementarity.^8^ A similar pattern emerged in our simulations: at a GFE threshold of −1.2 kcal/mol, positively charged FragMaps were widespread across the nucleic acid surface. While the C36 FF produced discrete and scattered positive FragMaps the Drude FF yielded continuous, favorable positive FragMaps along the DNA hairpirn backbone.(Supplementary Figure S2).

Although favorable interactions between oppositely charged species are expected, accu-rately modeling species of like charge poses a greater challenge for nonpolarizable FFs due to electrostatic repulsion. To further assess the Drude FF’s ability to capture such effects, we examined negative FragMaps obtained from the sampling behavior of the negatively charged solute acetate. In the DDX simulations, negative FragMaps are present along the major groove, particularly near N4 of Cyt6, N6 of Ade8, N2 of Gua5, and N1 of Gua10 (Figure 2). With the C36 FF, the negative FragMaps were confined to two distinct sites, whereas with the Drude FF, they were more widely distributed throughout the groove. For nucleic acids with more complex tertiary structures, such as TAR and IVP, the major grooves are more tightly enclosed by the backbone, thus limiting acetate access to those regions due to the closer proximity of the negatively charged phosphate groups. Consequently, negative FragMaps in these systems were primarily localized near exposed nitrogen atoms of terminal or loop residues (Supporting Figures S4 and S5).

The presence of cations also influenced acetate sampling. Systems containing K^+^ (TGQ) or Mg^2+^ (HCV and GUA) exhibited extensive acetate localization near the ions (Supplementary Figures S3, S5, and S7). In contrast, the previous SILCS-RNA study reported minimal acetate sampling around Mg^2+^, a behavior consistent across most C36 systems except HCV, where negative FragMaps were present within the minor groove and is a result reproduced in this study (Figure 3).^8^ Solute sampling around cations is primarily governed by two factors: the solvent accessibility of the ion and its ability to diffuse from its initial position. These effects were most evident in Mg^2+^-containing systems, where acetate was predominantly localized near solvent exposed Mg^2+^ ions for both FFs. However, the Drude FF produced more symmetric interactions around the ions (Figure 3B). While Mg^2+^ ions in HCV showed limited diffusion, substantial positional shifts were observed for GUA, leading to sparse or absent acetate sampling that reflected ion mobility rather than reduced favorable interactions (Figure 3C).

**Figure 3:**
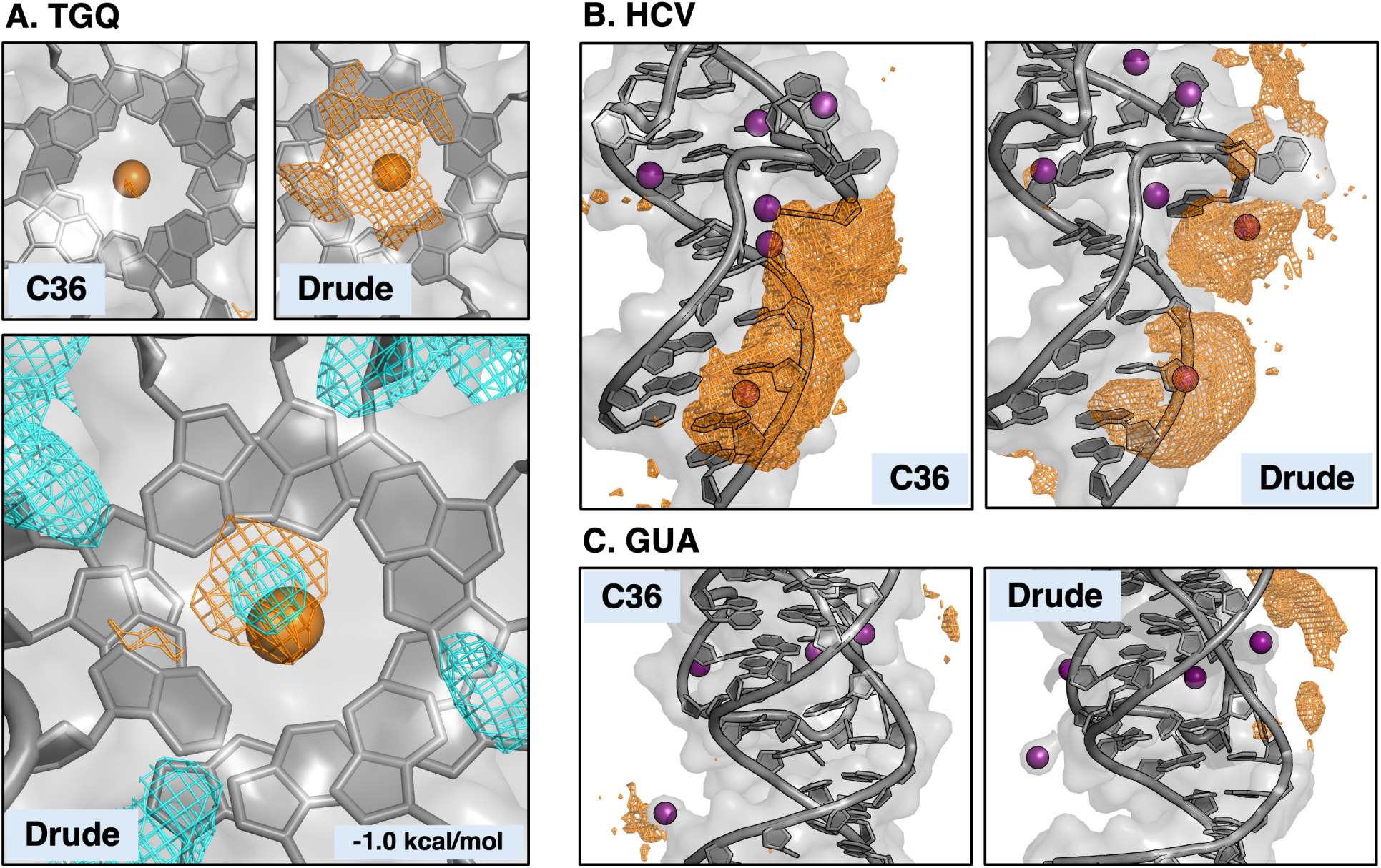
Acetate FragMaps for TGQ, HCV, and GUA. All ACEO (orange) maps are reported at −0.5 kcal/mol GFE unless labeled otherwise. (A) TGQ occupancy maps surrounding the GQ core. A comparison (bottom) of ACEO (orange) and MAMY (cyan) indicate charged occupancy overlap above the K^+^ ions and the electronegative core, at the same GFE level. (B,C) HCV and GUA acetate FragMaps around exposed Mg^2+^ ions.

Evaluating how internal K^+^ ions, such as those in GQ structures, influence binding site properties is crucial for improving ligand design. These cations reside within the electroneg-ative channel formed by the carbonyl oxygen atoms of the guanine tetrads, where they play an essential role in the folding and structural stability of GQ. Polarizable FFs like Drude have been shown to more accurately model these ion-nucleic acid interactions.^31,32,58^ With the Drude FF, acetate is localized above the positively charged K^+^cations in the GQ core, whereas no such sampling was observed with C36. (Figure 3A, top). While the localization likely reflects electrostatic attraction to the core cations, guanine nitrogen atoms may also contribute to acetate coordination above the core, similar to their enrichment around major groove-facing nitrogens in DDX (Figure 2. This region is a known compound binding site in GQs, typically exploited by ligands through *π*-*π* interactions or cationic interactions aligned with the core ions. Notably, the concurrent localization of methylammonium and acetate at similar GFE thresholds (Figure 3A, bottom) suggests that negatively charged ligand moieties might also engage this site, presenting a new avenue for GQ targeted ligand design.

Altogether, the enhanced acetate localization near cations with the Drude FF demonstrates that incorporating electronic polarization increases sampling of negatively charged solutes around polyanionic nucleci acids compared to nonpolarizable FFs like C36. This observation aligns with previous reports showing improved modeling of ion condensation around DNA using the Drude FF.^59,60^ Moreover, our findings suggest that including polarization not only resolves some limitations of the original SILCS-RNA approach^8^ but also enables new hypotheses for targeting nucleic acid systems.

To further investigate the FragMap differences observed between the C36 and Drude FFs, we analyzed the molecular dipole moments of the eight SILCS solute molecules. We hypothesized that the ability of small molecules to adapt their electronic structure to different microenvironments, as a function of proximity to the highly charged nucleic acids, contributed to the subtle differences in solute sampling. Solute dipole moments were calculated from the Drude SILCS simulations to assess how each solute’s electronic response varied across the nucleic acid structures.

The dipole moment distributions of the SILCS solutes spanned roughly 1 - 2 D (Figure 4), reflecting fluctuations in solute polarization arising from changes in position, orientation, and local interactions. These distributions were consistent for each solute across all nucleic acid systems, indicating that dipole variability is an intrinsic feature of the molecules in the presence of nucleic acids and rather than a feature of specificc nucleic acid topologies. This sensitivity to the surrounding electric field likely contributes to the subtle differences observed in the Drude FragMaps. As electronic polarization is inherently directional, with certain hydrogen-bonding groups treated anisotropically through the use of lone pairs and atomic anisotropic polarizability, the Drude model allows for more sensitive responses to such electrostatic effects.^61^

**Figure 4:**
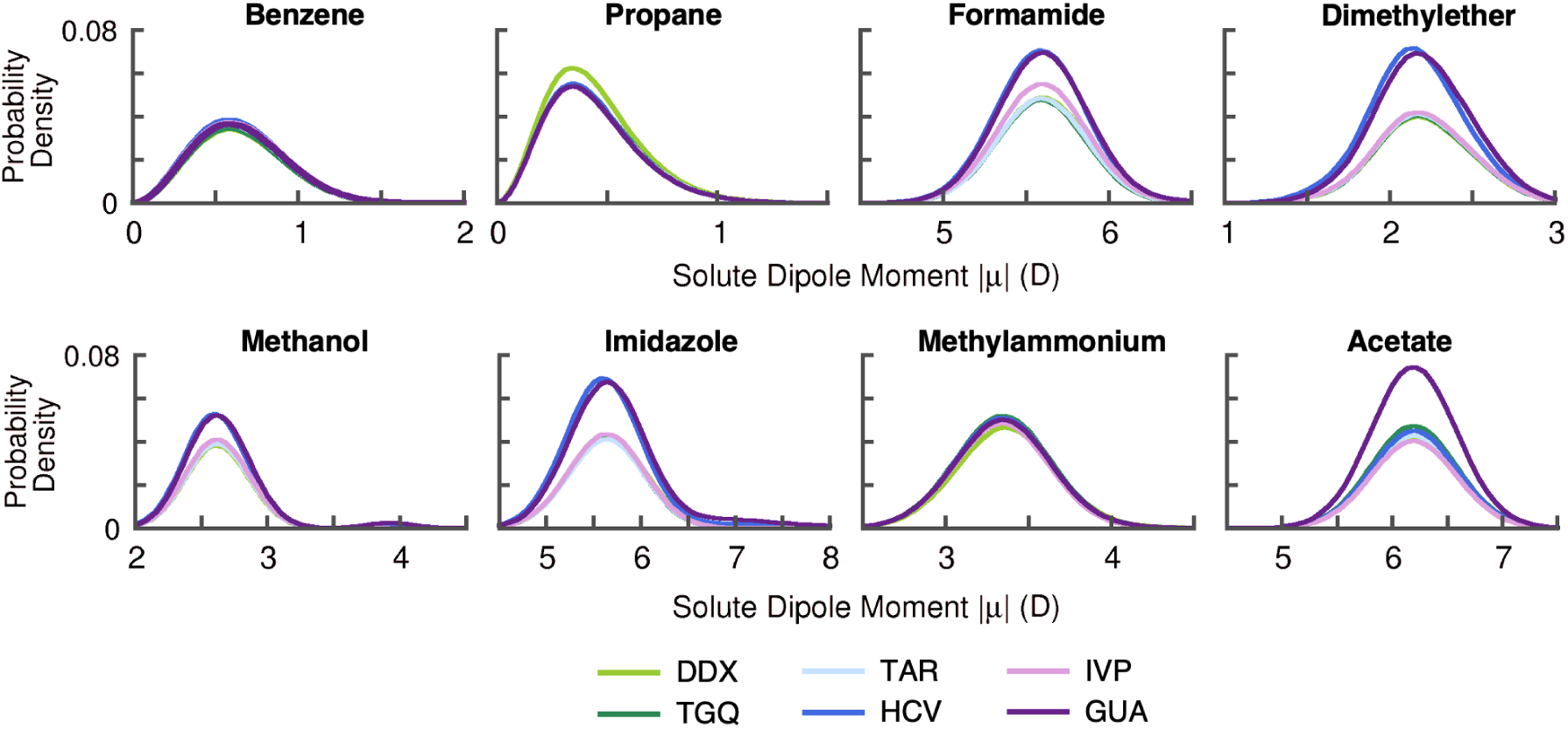
Distributions of solute dipole moments for all Drude systems.

This improved sensitivity also impacts the spatial distribution of each solute relative to the nucleic acid structures. Neutral solutes displayed two preferred localization regions, at approximately 3 Å and 15 Å from the the nucleic acid surface, indicating both short- and long-range accumulation (Figure 5). As expected, the charged solutes exhibited distinct behaviors: positively charged methylammonium primarily localized within 3 Å of the negatively charged phosphate backbone, while acetate was predominantly found farther away, with a major peak near 15 Å due to electrostatic repulsion. Small secondary peaks in the ac-etate distributions near 3 Ålikely reflect interactions with nearby cations and solvent-exposed nucleobase nitrogens consistent with the localization patterns discussed above.

**Figure 5:**
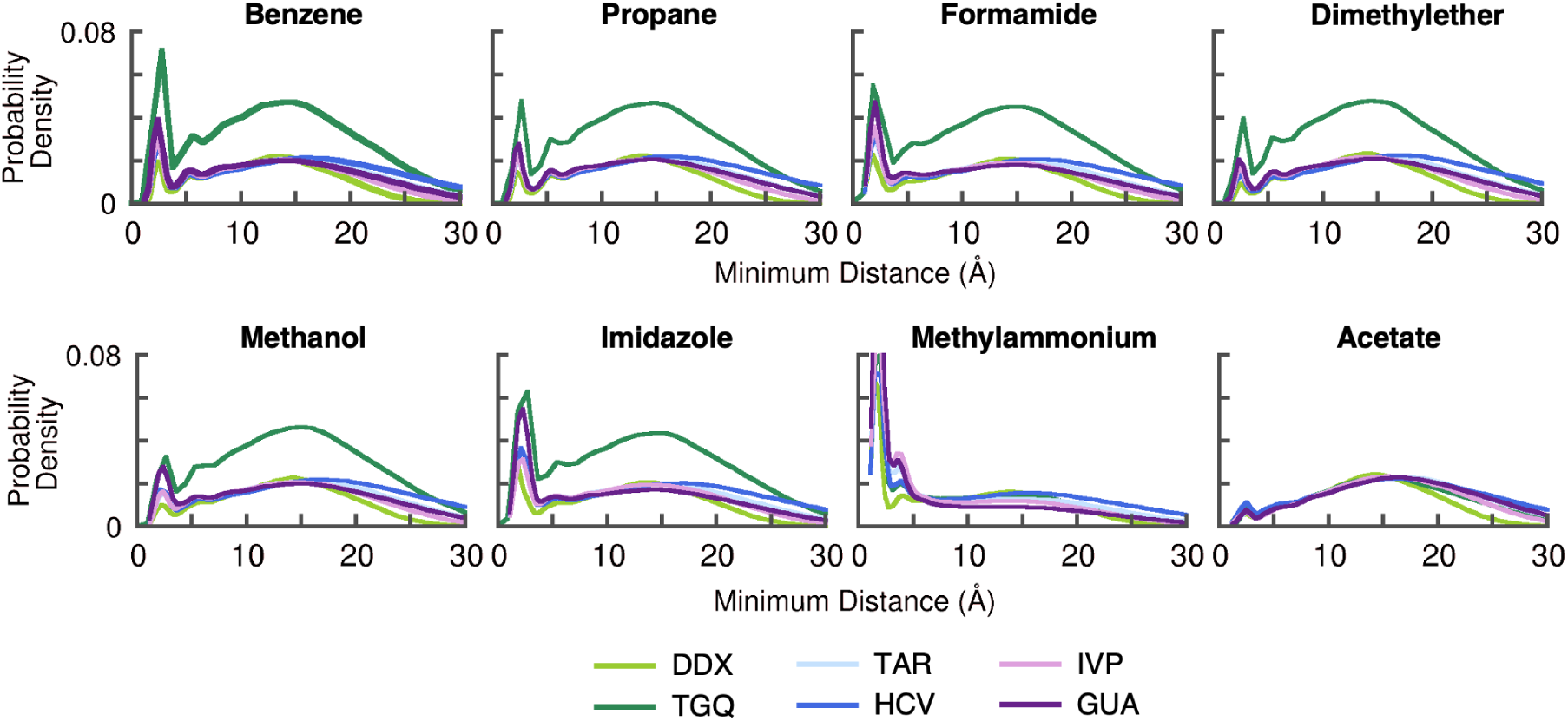
Distributions of solute minimum distance from the biomolecule in all Drude sys-tems.

Together, analysis of molecular dipole moments and localization suggests that the SILCS solutes are electronically influenced by the DNA and RNA targets. Their molecular dipole moments exhibit broad ranges and each solute shows characteristic accumulations both close to the nucleic acid surfaces and at greater distance. The perturbations in dipole moments due to changes in electronic structure are a feature only observable with polarizable FFs, and are comparable to the previous report of polarizable SILCS in the context of proteins.^8^ As a result, combined with the differences between C36 and Drude FragMaps, our results suggest that there are subtle but important features that are not captured by nonpolarizable FFs that can be observed with the Drude FF. We expand on the importance of this concept in the following sections.

### Identification of Known Binding Sites

Accurately identifying known binding sites and binding interactions is a crucial benchmark for new drug discovery methodologies. In the case of SILCS, the ability of the simulations to accurately map favorable binding locations of known ligands would verify the use of SILCS to identify novel binding sites. With the exception of DDX, all nucleic acid structures studied here were experimentally resolved as ligand-bound structures with different classes of interactions such as major and minor groove binders, electrostatic surface binders, pocket binders, and intercalators. While our validation set does not incorporate every type of nucleic acid - small molecule interaction, it serves as a robust test of the ability of SILCS to identify a diverse set of binding modes commonly seen in small molecules that target nucleic acids.

### Major and Minor Groove Binding Sites

IVP forms a partial duplex that is selectively recognized by an RNA-dependent RNA polymerase (RDRP) for regulation of Infleunza A viral transcription. In attempts to modulate transcription, a novel anti-influenzae ligand scaffold, 6,7-dimethoxy-2-(1-piperazynl)-4-quinazolinamine (DPQ) was identified as a major groove binder.^28^ Despite its solvent accessibility, the major groove in IVP had limited solute sampling using both FFs (Figure 6). Only acetate, methanol, and apolar solute sampling (although present at an unfavorable GFE threshold) was observed with C36, agreeing with the previous SILCS-RNA study. In contrast, the Drude SILCS-Nucleic approach lead to more sampling of apolar and hydrogen bond donor and acceptors yieliding more favorable GFE FragMaps. Unsurprisingly, the limited sampling resulted in no C36 FragMap overlap with DPQ, while Drude donor and acceptor maps were shown to overlay with DPQ near a dimethoxy group and the piperazynl ring (Figure 6). The sparse favorable solute sampling at this site may help explain the weak binding affinity of DPQ to IVP^28^ but also raises the question of can the major groove be effectively targeted with small molecules to out-compete RDRP. The lack of C36 FragMap density in a ligand binding site identified experimentally is concerning, but the observed improvement with Drude SILCS-Nucleic suggests that the polarizable FF approach better models this solvent-accessible site.

**Figure 6:**
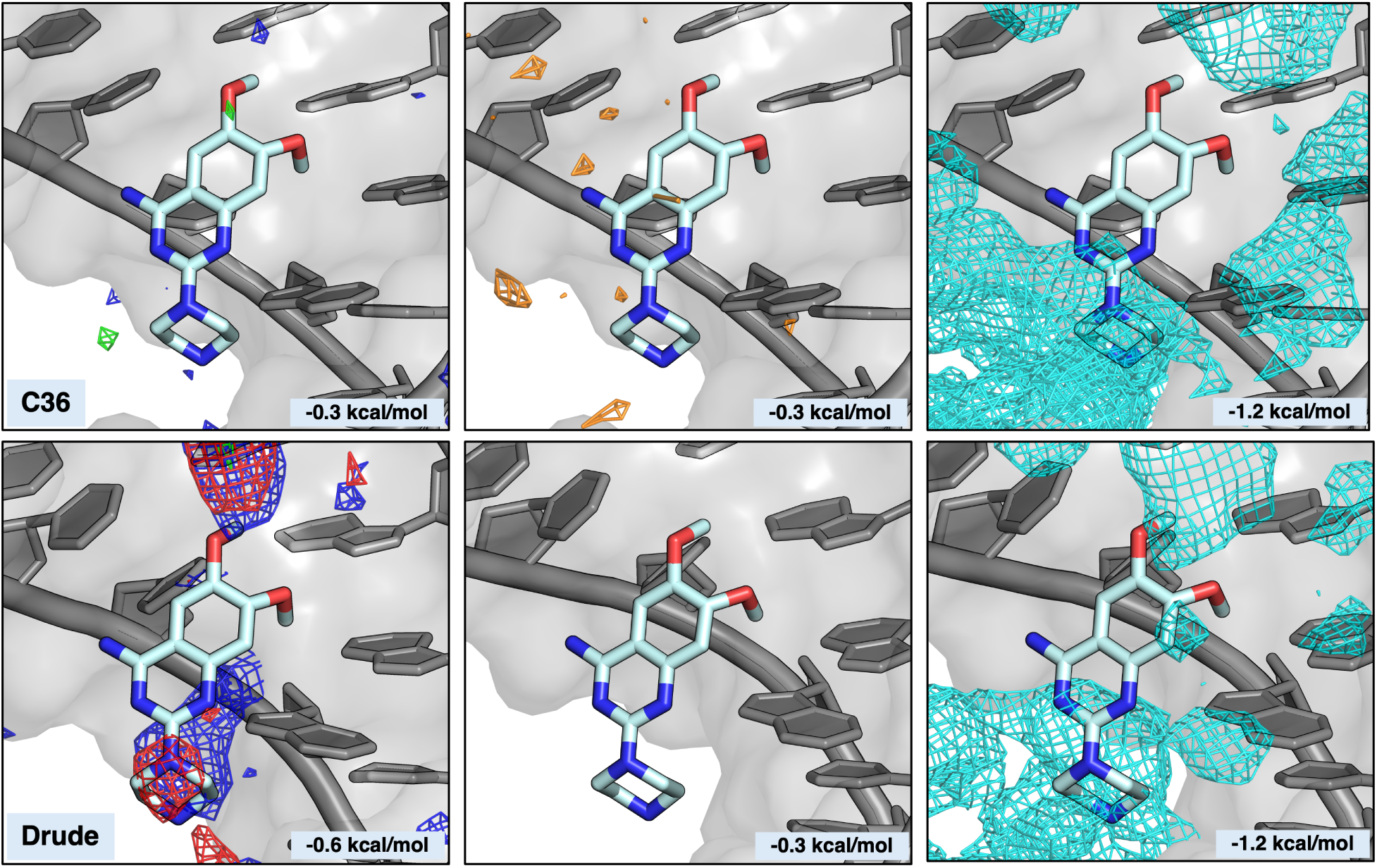
Binding site interactions for IVP with the ligand DPQ. FragMaps are shown as apolar (light green), hydrogen bond donor (blue), hydrogen bond acc (red), negatively charged (orange), and positively charged (cyan). GFE thresholds are given in each panel.

In the case of HIV-1 TAR RNA, both proteins and small molecules have been shown to bind in the major groove for activation and inhibition of the proviral genome.^26,62–64^ The chosen TAR structure (PDB: 6CMN) was co-crystallized with a designer RNA recognition motif (RRM) that interacts with TAR via hydrogen bonds and salt bridges between arginines in the RRM and three guanine bases in TAR. Guanidinium moieties are commonly found in small molecules designed to bind nucleic acids to form favorable interactions with the negatively charged phosphate backbone that are reminiscent of arginine.^65^ Methylammonium sampled the major groove with both the C36 and Drude FFs is evident, particularly at sites that coincide with the arginines in the designer RRM. The small molecule RBT203, a bis-guanidine compound that also interacts with a slightly different TAR loop conformation through its positively charged guanidinium groups, yielded similar overlap with positively charged maps (Figure S8).^63^

Within the minor groove of TAR, Neomycin B (NEOB) has been shown to bind and prompt a conformational change in the RNA structure. While the conformation used in our SILCS simulations is not the same as one that was resolved when bound to NEOB (PDB: 1QD3),^62^ we were interested to see if the Drude FF could better identify its binding site on this structure compared to C36. The previous SILCS-RNA study reported solute sampling in the minor groove at a weak GFE threshold of −0.4 kcal/mol. Here, we obtained similar results showing sparse solute sampling with C36 and a GFE threshold of −0.6 kcal/mol (Figure 7). In contrast, solute sampling in the minor groove was observed at a much more favorable GFE threshold of −1.0 kcal/mol for the Drude FragMaps, reinforcing the findings from DDX. That is, the Drude FF yields more specific interactions within minor grooves. The lack of FragMap occupancy in the minor groove with the C36 FF mirrors that of the IVP major groove and suggests a weakness in the ability of the additive FF to predict such binding sites. The polarizable FF was better able to capture these interactions.

**Figure 7:**
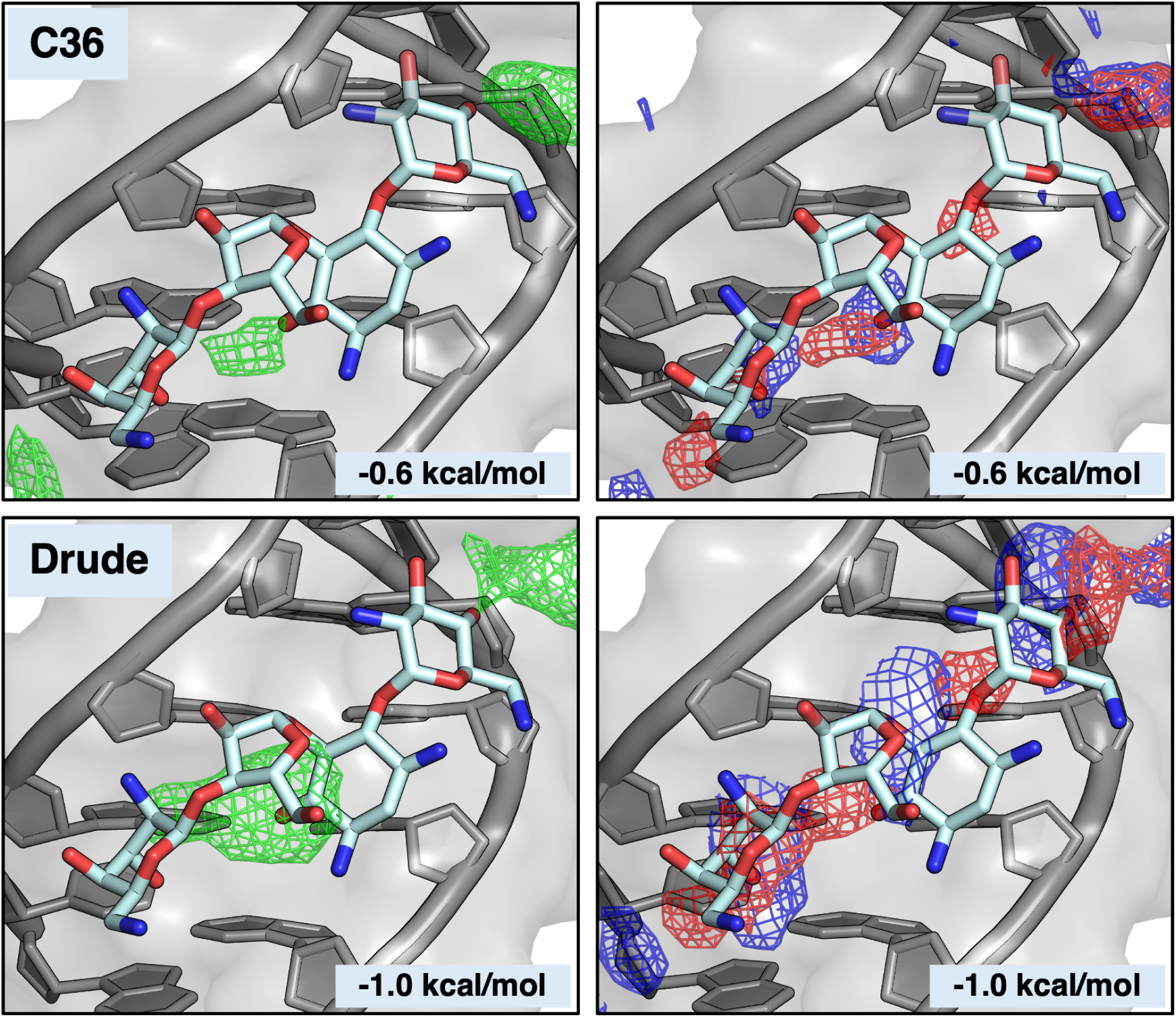
NEOB binding interactions within the TAR minor groove. Apolar (light green) and hydrogen bond donor (blue) and acceptor (red) sampling was observed on the face of the groove.

### Structural pocket binders and Intercalators

The internal ribosome entry site in the hepatitis C virus is responsible for the initiation of viral protein synthesis by recruiting 40S ribosomal subunits. Small molecule drugs such as benzimidazole (BENZ) and its derivatives are known to intercalate into this binding region.^27^ These compounds are typically aromatic and planar, thus their insertion is generally driven by apolar and aromatic stacking interactions between the compound and nearby bases (Figure 8A, top). The previous SILCS-RNA study reported apolar occupancy overlapping with BENZ only at GFE values below −0.5 kcal/mol. This was similar to our C36 results where no apolar occupancy was seen at −1.0 kcal/mol. In contrast, Drude simulations produced apolar occupancy at −1.0 kcal/mol, indicating a much more favorable interaction (Figure 8A, bottom). With the surrounding nucleobases, it was expected that there would be some favorable apolar sampling within this site that would overlap with the aromatic rings of BENZ with both FFs. The chemical nature of BENZ prompted us to investigate the FragMaps for benzene, propane, and imidazole to further elucidate the dominant type of interaction within the pocket. Imidazole FragMaps are generated for nitrogen atoms (hydrogen bond donors and acceptors) and for the apolar carbon atoms. Therefore, imidazole sampling can be monitored for the balance of polar and apolar interactions. Both propane and imidazole sampled the binding pocket, whereas benzene was completely absent from the pocket with both FFs despite sampling around nucleobases elsewhere in the structure (Supplementary Figure S9). It is therefore likely that solutes containing hydrogen bond donor groups out-compete the larger benzene solute as the generic hydrogen bond donor and acceptor maps (to which imidazole contributes) are predominantly located in and around the binding pocket. The binding pocket is defined by nucleobases, such as the known interactions between BENZ and nearby Gua110, which may act to stabilize the BENZ binding pose. This observation may also explain the increase in apolar sampling with Drude as it contains contributions of the imidazole C atoms within the binding site. The hydrogen bond donor and acceptor FragMaps were more favorable within the binding pocket at −1.2 and −1.4 kcal/mol for C36 and Drude, respectfully. This observation provides further evidence that these interactions dominant ligand binding in the pocket.

**Figure 8:**
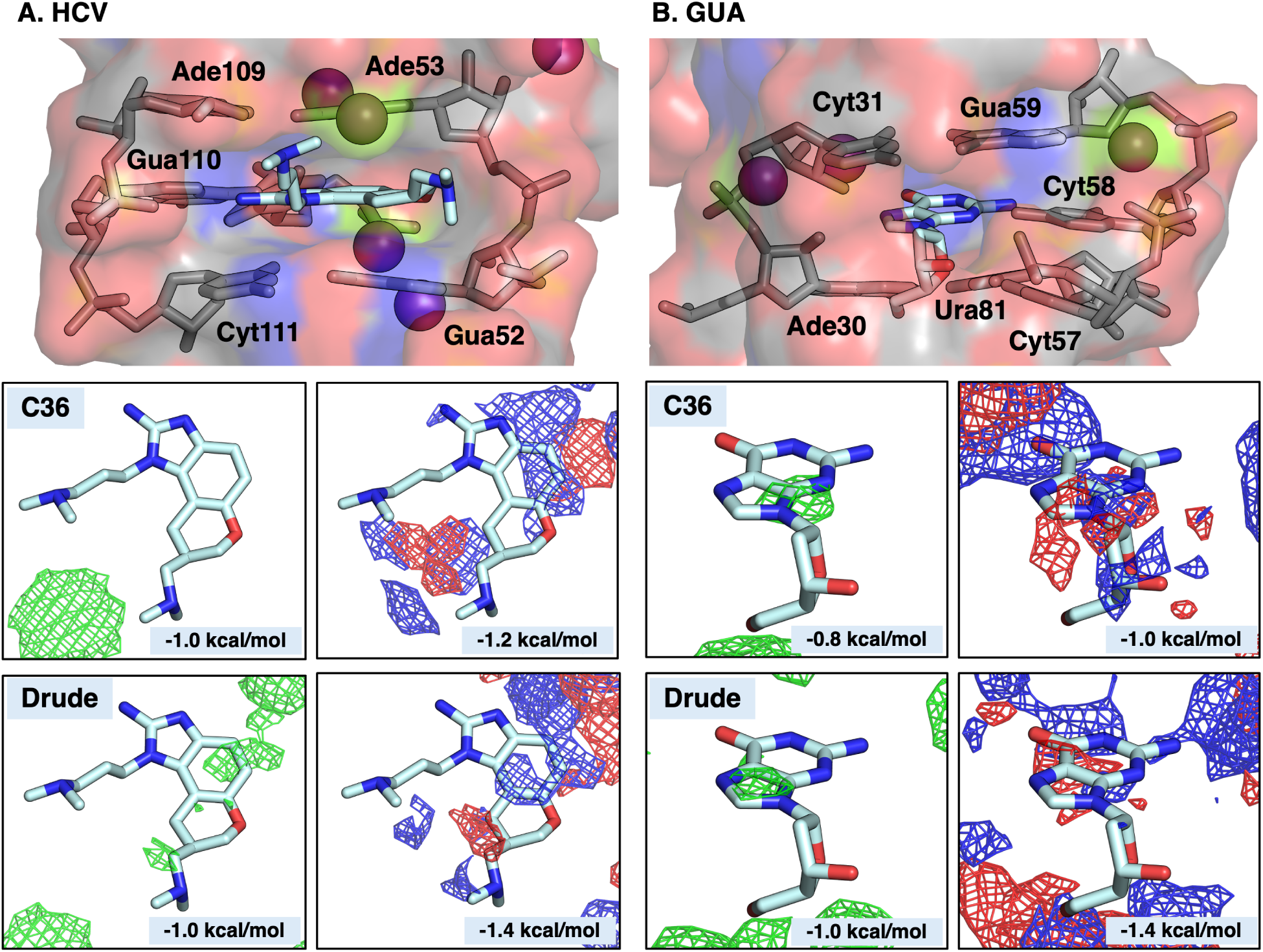
HCV and GUA binding interactions. (A) BENZ intercalates into the HCV major groove near Mg^2+^. (B) dG binds within the GUA structure and is nearly entirely occluded from solvent. Both binding locations are shown as surface and colored by atom type with potential interacting residues shown as sticks.

Riboswitches contain binding pockets that are highly specific for their effector molecules, which can provide insight into modifications that can be exploited for antimicrobial agents.^29^ The binding site of GUA for deoxyguanosine (dG) is nearly inaccessible within the structure but was sampled by solutes using both C36 and Drude (Figure 8A, top). As with the other systems, the Drude FragMaps were systematically more favorable than those of C36. Apolar maps were localized near the junction of the pyrimidine and imidazole rings in C36 and directly overlapped with the imidazole ring with Drude. Hydrogen bond donor and acceptor maps were observed to overlap nitrogen and oxygen atoms on dG, corresponding to sites of hydrogen bonding interactions with the surrounding nucleobases in the pocket (Figure 8B, bottom). This outcome may help to explain the high selectivity of GUA for dG as a result of the complementarity between the structural and chemical characteristics of the binding pocket and the functional groups of dG.

### Surface binders

Some noncanonical nucleic acid structures have distinct structural fea-tures that have primarily been the focus for ligand binding. Specifically for GQ structures, ligands primarily interact via *π* − *π* stacking interactions on the surface of exposed guanine tetrads. As such, these ligands typically contain a central aromatic core with additional functional groups to interact with other parts of the GQ structure, such as the backbone or loops. Napthalene diimide (NDI) is a common scaffold for GQ ligands and the NDI derivative MM41 is known to stabilize TGQ, ultimately inhibiting cell proliferation in pancreatic cancer cells.^25^ MM41 is a symmetrical ligand with arms containing *N* -methylpiperazine and morpholine groups that can interact with the TGQ grooves, yielding a distinct interaction fingerprint for SILCS to replicate within a binding site exposed to all solutes.

Both FFs produced apolar FragMaps that overlapped with the NDI core of MM41 (Figure 9), which agrees with the known *π*-*π* interactions involving tetrad guanines. In addition, the arms of MM41 are hypothesized to interact with the phosphate backbone via hydrogen bonding and charge-charge interactions. The favorable hydrogen bond donor and acceptor Fragmaps with the Drude FF improved overlap with nitrogen and oxygen atoms located in the arms of MM41 compared to C36. We also observed positive FragMaps near the TGQ grooves that has some overlap with the protonated *N* -methylpiperazine and morphilino nitrogens on two of the arms. Thus, in the case of TGQ, whereas the C36 and Drude FFs largely produce the same features that characterize the NDI core of MM41, the Drude FF yields a better description of the flexible arms that involve polar and charged interactions with the phosphodiester backbone and chemical moieties in the grooves.

**Figure 9:**
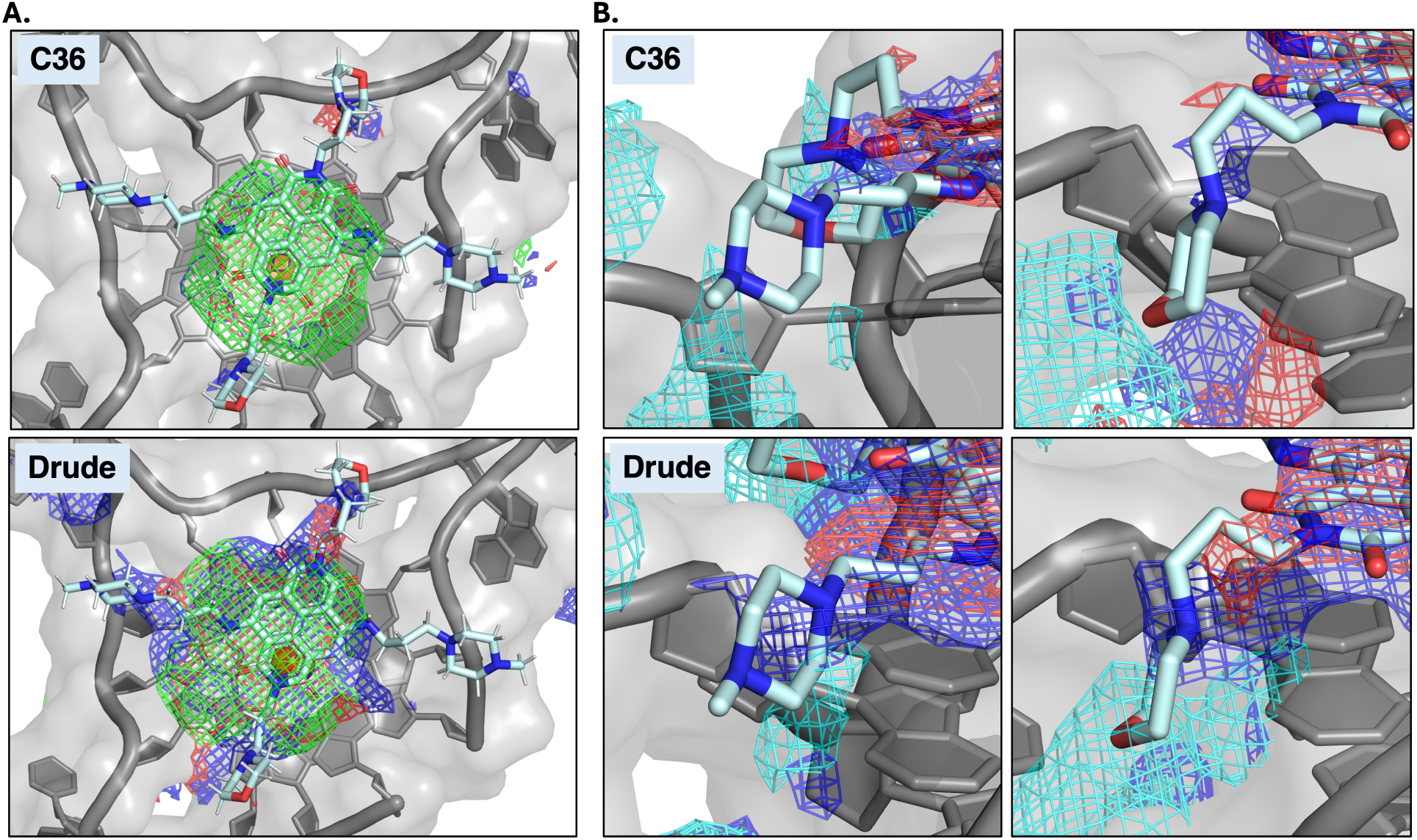
FragMap overlap with MM41. A) Apolar (light green) and hydrogen bond donor (blue) and acceptor (red) maps are visualized around the GQ core at GFE levels of −1.0 kcal/mol and −0.8 kcal/mol for C36 and −1.0 kcal/mol and −1.0 kcal/mol for Drude. B) Positively charged maps (cyan) and hydrogen bond donor (blue) and acceptor (red) maps were observed near N-methylpiperzine (left) and morphilino (right) groups that are hypothesized to be protonated and form hydrogen bonds with the GQ backbone. Positvely charged maps are shown at −1.5 and −1.8 kcal/mol for C36 and Drude. Hydrogen bond donor and acceptor maps are shown at −0.6 and −0.8 kcal/mol for C36 and Drude, respectively.

### Comparison to Molecular Docking

An essential aspect of computational drug discovery is the ability to not only identify binding sites but also present hypotheses for the best ligand poses within those binding sites. Molecular docking approaches have been traditionally used to identify ligand poses but rely on static target structures and limited electrostatic considerations for nucleic acids. As such, we wanted to determine if the internal dynamics and competition amoung solutes and water that are incorporated into the SILCS FragMaps, along with the improved consideration of electronic properties with the Drude FF, could better replicate known experimental poses of ligands to the studied nucleic acid structures compared to popular docking methods such as Schrodinger Glide and Gnina.

Within the SILCS software suite, SILCS-MC can be used to refine ligand conformations based on overlap of ligands with the FragMaps yielding the LGFE scores. SILCS-MC uses a spherical region for ligand sampling compared to the standard box used in both Glide and Gnina. The same x-, y-, and z-center of the sampling region remained the same for each method while box and sphere sizing was adjusted due to differences in the ligand sampling methodologies. At maximum, the top 9 poses were obtained from each method along with scores unique to each method that have components to predict the strength of binding and evaluate each pose conformation. For SILCS-MC, docking is guided primarily by the LGFE, the SILCS metric for binding affinity. This value can be normalized across compounds by dividing by the number of atoms within the ligand to calculate a ligand efficiency (LE). Glide produces a docking score that determines the strength of binding; however, a more robust method that incorporates solvation, such as MM-GBSA, is typically used to obtain a more accurate prediction of binding affinity. Gnina also has its own approach to determine ligand affinity including the intramolecular energy of the ligand to quantify the energy penalty for the specific pose. These values along with each poses RMSD are reported in Supplemental Tables S8, S9, S10.

RMSD was used to determine the extent of deviation between the generated poses from each method and the experimental crystal pose. For all groove binding ligands, SILCS-MC resulted in the lowest RMSD poses (all under 3 Å), whereas both Glide and Gnina resulted in higher RMSD values (Table 2). By overlapping the top poses with the crystal structure, the SILCS-MC poses were shown to better replicate the position of core ring structures in DPQ, RBT203, and NEOB compared to Glide and Gnina. RRM was excluded from pose visualization images as both Gnina and Glide wrapped the peptide around the major groove rather than maintaining the location of the arginine residues, which explains the higher RMSD for RRM with these methods despite RMSD only being calculated for the arginine residues.

**Table 2:** Lowest RMSD (Å) values for groove binding ligands using C36 and Drude structures from SILCS simulations.

|  | **DPQ** |  | **RBT203** |  | **RRM** |  | **NEOB** |  |
| --- | --- | --- | --- | --- | --- | --- | --- | --- |
|  | C36 | Drude | C36 | Drude | C36 | Drude | C36 | Drude |
| SILCS-MC | **2.64** | **2.19** | **2.96** | **2.50** | **1.55** | **1.49** | **2.68** | **2.10** |
| Glide | 4.36 | 4.69 | 6.76 | 8.94 | 8.65 | 12.9 | 5.13 | 6.03 |
| Gnina | 4.61 | 4.52 | 8.85 | 4.63 | 8.29 | 7.71 | 3.67 | 3.72 |

**Table 3:** Lowest RMSD (Å) values for pocket binding, surface binding and intercalating ligands.

|  | **dG** |  | **BENZ** |  | **MM41** |  |
| --- | --- | --- | --- | --- | --- | --- |
|  | C36 | Drude | C36 | Drude | C36 | Drude |
| SILCS-MC | **1.11** | **0.847** | 2.90 | 1.89 | 8.75 | **4.24** |
| Glide | 10.7 | 0.993 | 1.58 | 1.74 | **8.51** | 6.35 |
| Gnina | 3.19 | 7.75 | **0.788** | **1.15** | 11.8 | 6.12 |

SILCS-MC was also able to best replicate the binding of dG to GUA, with an RMSD under 1.5 Å for both C36 and Drude despite the pocket being almost entirely occluded from solvent. However, Gnina outperformed both SILCS-MC and Glide for the intercalating compound, BENZ, but SILCS-MC still obtained a RMSD less than 3 Å, which was similar to both Glide and Gnina. The main differences between the BENZ poses occurred in the orientation of the compound within the pocket, which could be due to the lack of apolar sampling during both C36 and Drude simulations (Supporting Figure S11B). The highest RMSD values produced by SILCS-MC were observed for the GQ surface binding compound, MM41, although Glide only outperformed SILCS-MC using the C36 structures by less than 0.2 Å. The generated poses all resulted in differences in the positions of the flexible arms containing the *N* -methylpiperazine and morpholino groups, which are hypothesized to interact within the backbone grooves based on the experimental structure (Supporting Figures S11C).

The improved performance of SILCS-MC using the Drude-generated FragMaps highlights how subtle differences in solute sampling can have a substantial impact on accurately predicting ligand binding. Across all ligands the Drude FragMaps consistently produced poses with lower RMSDs, compared to C36. These results reflect the cumulative effect of improved treatment of electrostatic interactions and how these subtleties are overlooked in other docking methods such as Glide or Gnina. Taken together, these findings emphasize that accurately modeling nucleic-acid interactions requires both consideration of dynamics and polarization response and that SILCS-Nucleic with the Drude FF provides a clear advantage over popular docking approaches.

## Conclusions

Presented here is an extension of the SILCS-RNA workflow that incorporates the Drude polarizable FF to allow explicit treatment of electronic polarization effects during ligand-nucleic acid interactions. As anticipated, the inclusion of electronic polarization effects improved the modeling of ligand-nucleic acid interactions. FragMaps generated with the Drude FF showed enhanced solute localization, orientation, and interaction favorability, leading to more accurate identification of binding sites across diverse nucleic acid structures. Notably, the Drude FF also resolved key shortcomings of the previous SILCS-RNA study, as negatively charged acetate solutes sampled regions near Mg^2+^ ions and other strongly polarized environments. This behavior reflects the ability of the Drude model to capture inducible dipole fluctuations that fixed-charge FFs fundamentally cannot represent.

Our results highlight a broader challenge in the field. Most common computer-aided drug design tools were developed for protein targets and may not account for the electrostatic, structural, and dynamical properties of RNA and DNA that differ from proteins. Docking programs such as Glide and Gnina, for example, were built using scoring and sampling strategies optimized for protein environments and interactions, which may underlie their struggles with the polyanionic backbone, flexible grooves, and unconventional binding modes common in nucleic acids. In our study, SILCS-MC produced the most accurate groove-binding poses, outperforming both Glide and Gnina. These traditional docking methods, however, were competitive for pocket-binding, intercalating, and surface-binding ligands, emphasizing that no current approach fully addresses the challenges of modeling nucleic acid-small molecule interactions.

Taken together, this work demonstrates that incorporating electronic polarization into a computational workflow can close the gap between current methods and the biophysical nature of nucleic acids. As interest in RNA- and DNA-targeted therapeutics continues to grow, especially with the continued understanding of noncoding RNA biology and RNA-modulating small molecules, such approaches offer a promising path forward. Continued development of these methods, alongside benchmarking across diverse nucleic acid motifs and ligand classes, will be essential for advancing computational strategies that can reliably support nucleic acid-focused drug discovery.

## Author Contributions

HMM: Conceptualization, Investigation, Methodology, Formal analysis, Writing - original draft, Writing - review and editing

AMB: Investigation, Methodology, Writing - review and editing

ADM: Methodology, Writing - review and editing, Funding acquisition

JAL: Conceptualization, Methodology, Writing - review and editing, Project administration, Funding acquisition

## Data Availability

All code necessary to reproduce the findings presented here and to implement the Drude SILCS-Nucleic workflow for other systems is available at https://github.com/hmmichel/drude_silcs_nucleic.

## Supplementary Data

Supplementary Data are available at NAR Online.

## Acknowledgements

The authors thank Virginia Tech Advanced Research Computing and the University of Maryland Computer-Aided Drug Design Center for computing time and resources, and Dr. Abhishek Kognole for helpful discussions. This work also used the Expanse Cluster at San Diego Supercomputer Center (SDSC) through allocation BIO230117 (to JAL) from the Advanced Cyberinfrastructure Coordination Ecosystem: Services & Support (ACCESS) program, which is supported by National Science Foundation grants #2138259, #2138286, #2138307, #2137603, and #2138296.

