## Supplementary Information for "Drude SILCS-Nucleic: Harnessing Explicit Electronic Polarization in Targeting RNA and DNA for Drug Design"

**Methods S1.**

**Formamide Parametrization for the Drude FF.** In order to apply the Drude SILCS-Nucleic process to nucleic acids, we needed to evaluate and improve the Drude parameters for formamide. Seven formamide models were generated with FFParam using different target data, such as fitting bonded terms to quantum mechanical (QM) vibrational frequency or targeting dimer or trimer dipole moments and molecular polarizabilities. All QM geometry optimizations and vibrational frequency calculations were performed in Gaussian09[1] using an MP2/6-13+G* model chemistry. All dipole moment and molecular polarizability calculations were performed in Psi4[2] using an RIMP2/cc-pVQZ model chemistry.

Electrostatic parameters were adjusted with different windows on amide hydrogen atoms or by allowing adjustments to anisotropy terms around specified atoms, including the nitrogen atom of the amide group. All models were evaluated by their ability to replicate experimental heat of vaporization ($\Delta H_{vap}$), dielectric constant ($\varepsilon$), and density, allowing us to evaluate the strength of the bonded terms and electrostatic parameters. To obtain these values, the CHARMM program[3] was used to generate pure formamide systems in liquid phase, containing 216 formamide molecules, at 298 K. Monomer simulations were also conducted with a single molecule in the gas phase. $\Delta H_{vap}$ was then determined according to Eq. 1.

$$\Delta H_{vap} = \langle U_{gas} \rangle - \frac{\langle U_{liq} \rangle}{N_{mol}} + RT \tag{1}$$

where $\langle U_{gas} \rangle$ and $\langle U_{liq} \rangle$ represent the average potential energies of formamide in the gas and liquid phases, respectively. $N_{mol}$ corresponds to the number of molecules in the liquid simulation, in this case 216. $RT$ corresponds to the thermal energy at $T = 298\ K$.

Seven formamide models were generated and evaluated using a variety of parameter fitting schemes to determine their ability to model non-bonded interactions and replicate experimental $\Delta H_{vap}$ and $\varepsilon$ (Supplementary Figure S2). Ultimately, model 6 was chosen as it produced a $\Delta H_{vap}$ of 15.4 kcal/mol, which was was within the range of the experimental $\Delta H_{vap}$, 14.0-15.5 kcal/mol.[4,5] Model 6 was also in the top three in its ability to replicate $\varepsilon$, with a value of 96.8 D compared to 111 D experimentally,[6] and density, which was also slightly overestimated, but acceptable at 1.21 g/mol compared to 1.13 g/mol experimentally[7] (Supplementary Figure S2). The improvement in the ability of this model to replicate $\Delta H_{vap}$ and $\varepsilon$, translates to improvements in intermolecular interactions which are especially important for ligand binding studies. The CHARMM-formatted topology and associated refined parameters for this model are provided in the Supplementary Information.

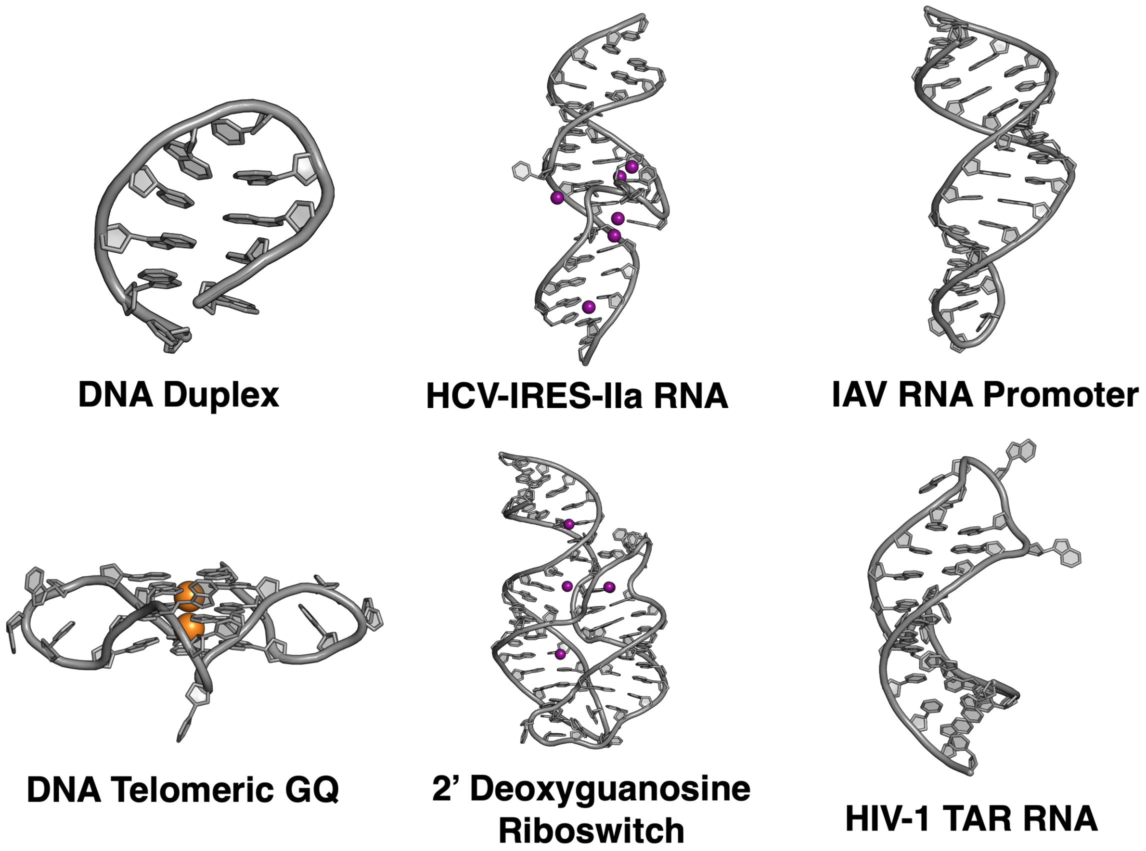


Figure S1: Structures of the DNA and RNA systems evaluated.

Table S1: SILCS atom types and associated FragMap types used for visualization

| Atom Type | Atom Location | Major Functional Group Represented | FragMap |
|---|---|---|---|
| **MEOO** | O on MEOH | HB donor and acceptor O atom from hydroxyls | MeoO |
| **IMIN** | N on IMIA/IMID | HB acceptor N atom in planar rings | HB Acc |
| **IMINH** | N(H) on IMIA/IMID | HB donor N atom in rings | HB Don |
| **FORO** | O on FORM | HB donor and acceptor O atom from hydroxyls | HB Acc<br>HB Don |
| **FORN** | N on FORM | HB donor N atom in amides | HB Don |
| **FORC** | C on FORM | Carbonyl C atom in aldehydes | – |
| **DMEO** | O on DMEE | HB acceptor O atom from ethers | HB Acc |
| **BENC** | 6 C on BENX | C atoms in aromatic ring | Apolar |
| **PRPC** | 3 C on PRPX | Aliphatic C atoms | Apolar |
| **GEHC** | 3 C on IMIA/IMID | Generic heterocycle carbon | Apolar |
| **MAMN** | N on MAMY | Positively charged N atom | Mamy |
| **MAMC** | C on MAMY | Conjugated C atom in cation | Mamy |
| **ACEO** | 2 O on ACEY | Negatively charged O atom in carboxylates, anions, etc. | AceO |
| **ACEC** | C on ACEY | Carbonyl C atom in carboxylates, P atom in phosphates, etc. | – |
| **TIPO SWM4O** | O on TIP3/SWM4 | O atom in water | Water |

Listing S1: CHARMM-formatted topology and parameters of the refined Drude polarizable model for formamide.

```
RESI FORM          0.000 ! formamide
!
!     Ha      Ht
!      \     /
!       C---N
!      //    \
!      O      Hc
!
GROUP
! fit6, jal 8/9/2023
ATOM  C      CD2O1C    0.428  ALPHA   -0.770  THOLE    0.838
ATOM  HA     HDP1C     0.053
ATOM  O      OD2C1D    0.000  ALPHA   -0.557  THOLE    1.238
ATOM  LPA    LPDO1    -0.365
ATOM  LPB    LPDO1    -0.197
ATOM  N      ND2A1    -0.476  ALPHA   -1.771  THOLE    1.217
ATOM  HC     HDP1A     0.296
ATOM  HT     HDP1A     0.261

BOND C    O    C    HA
BOND C    N    N    HC  N    HT
BOND O    LPA  O    LPB

LONEPAIR relative LPA O C HA distance 0.30 angle 91.00 dihe   0.00
LONEPAIR relative LPB O C HA distance 0.30 angle 91.00 dihe 180.00
! fit6
ANISOTROPY O C LPA LPB  A11 0.937 A22  1.151
ANISOTROPY N C HC  HT   A11 0.810 A22  1.063

IMPR C N O HA

IC O   C   N   HT        1.2250  125.00  180.00  123.50   1.0250
IC O   N   *C  HA        1.2250  125.00  180.00  114.00   0.0000
IC C   HT  *N  HC        1.3500  123.50  180.00  113.00   1.0250
IC HA  C    N  HC        1.3500  123.50    0.00  113.00   1.0250

read rtf card appe

BONDS
CD2O1C   ND2A1    420.00      1.370 ! FORM; silcs toppar; hmm/jal

DIHEDRALS
HDP1A    ND2A1    CD2O1C   HDP1C       2.500    2   180.00

IMPROPERS
! noting here that the term was removed
! improves agreement with QM vibrational frequencies
!ND2A1    HDP1A    HDP1A    CD2O1C      1.000    0      0.00 ! FORM, jal
```

Table S2: Comparison of different formamide models against indicated experimental data.

| Model | $\Delta H_{vap}$ (kcal/mol) | Exp. $\Delta H_{vap}$ | $\varepsilon$ | Exp. $\varepsilon$ | Density (g/mol) | Exp. Density | Molecular Volume ($Å^3$) | Notes |
|---|---|---|---|---|---|---|---|---|
| **Original** | 13.5 | **14.0-15.5 kcal/mol** | 105 | **111** | 1.20 | **1.13 g/mol** | 62.49 | Obtained from the Drude SILCS paper. [8] |
| **3** | 11.7 | | 69.4 | | 1.15 | | 64.84 | Bonded terms fit to QM vibrational frequency. |
| **4** | 13.7 | | 64.2 | | 1.19 | | 62.87 | Charge too symmetric on amide H. Tighter windows utilized for fitting charges. |
| **5** | 16.7 | | 72.3 | | 1.24 | | 60.32 | Dipole and molecular polarizability fit to dimer. |
| **6** | 15.4 | | 96.8 | | 1.21 | | 61.58 | Same as model 5 but the anisotropy constants around O and N were allowed to change. |
| **7** | 15.0 | | 90.1 | | 1.22 | | 61.53 | Adjusted C anisotropy. |
| **8** | 16.4 | | 101.6 | | 1.22 | | 61.17 | Dipole and molecular polarizability fit to optimized dimer geometries for both *trans* (1 H-bond) and *cis* (2 H-bonds) interactions between monomers. |
| **9** | 17.5 | | 106.7 | | 1.24 | | 60.28 | Dipole and molecular polarizability fit to trimer. |

Table S3: C36 and Drude Overlap Coefficients for all solutes in all systems

| Atom Type | DDX | | TGQ | | TAR | | HCV | | IVP | | GUA | | SILCS Type |
|---|---|---|---|---|---|---|---|---|---|---|---|---|---|
| | **C36** | **Drude** | **C36** | **Drude** | **C36** | **Drude** | **C36** | **Drude** | **C36** | **Drude** | **C36** | **Drude** | |
| **MEOO** | 0.794 | 0.840 | 0.783 | 0.836 | 0.788 | 0.839 | 0.857 | 0.942 | 0.791 | 0.839 | 0.758 | 0.802 | **MEOO** |
| **IMIN** | 0.798 | 0.831 | 0.787 | 0.832 | 0.784 | 0.829 | 0.785 | 0.895 | 0.786 | 0.827 | 0.751 | 0.775 | **IMIN** |
| **IMINH** | 0.797 | 0.830 | 0.787 | 0.832 | 0.784 | 0.828 | 0.788 | 0.898 | 0.785 | 0.826 | 0.750 | 0.780 | **IMINH** |
| **FORO** | 0.796 | 0.835 | 0.787 | 0.832 | 0.786 | 0.841 | 0.844 | 0.897 | 0.787 | 0.833 | 0.767 | 0.803 | **FORO** |
| **FORN** | 0.793 | 0.834 | 0.785 | 0.834 | 0.785 | 0.841 | 0.845 | 0.899 | 0.787 | 0.833 | 0.770 | 0.808 | **FORN** |
| **FORC** | 0.794 | 0.833 | 0.787 | 0.831 | 0.786 | 0.839 | 0.844 | 0.897 | 0.787 | 0.832 | 0.768 | 0.805 | **FORC** |
| **DMEO** | 0.793 | 0.842 | 0.787 | 0.834 | 0.787 | 0.837 | 0.875 | 0.966 | 0.787 | 0.833 | 0.779 | 0.826 | **DMEO** |
| **BENC** | 0.905 | 0.914 | 0.894 | 0.905 | 0.900 | 0.910 | 0.904 | 0.951 | 0.901 | 0.910 | 0.893 | 0.896 | **BENC** |
| **PRPC** | 0.884 | 0.900 | 0.870 | 0.895 | 0.872 | 0.894 | 0.896 | 0.912 | 0.871 | 0.891 | 0.877 | 0.892 | **PRPC** |
| **GEHC** | 0.877 | 0.882 | 0.867 | 0.883 | 0.868 | 0.882 | 0.852 | 0.949 | 0.869 | 0.882 | 0.833 | 0.836 | **GEHC** |
| **MAMN** | 0.818 | 0.829 | 0.824 | 0.840 | 0.827 | 0.859 | 0.816 | 0.838 | 0.832 | 0.878 | 0.801 | 0.833 | **MAMN** |
| **ACEC** | 0.789 | 0.826 | 0.780 | 0.822 | 0.788 | 0.825 | 0.837 | 0.878 | 0.788 | 0.827 | 0.788 | 0.823 | **ACEC** |
| **ACEO** | 0.847 | 0.872 | 0.841 | 0.869 | 0.847 | 0.872 | 0.882 | 0.921 | 0.847 | 0.899 | 0.846 | 0.869 | **ACEO** |
| **TIPO SWM4O** | 0.986 | 0.987 | 0.985 | 0.986 | 0.986 | 0.995 | 0.982 | 0.984 | 0.985 | 0.986 | 0.983 | 0.984 | **TIPO SWM4O** |

Table S4: Average and standard deviation of the difference in OC between C36 and Drude systems.

| Atom Type | Average ± Std. Dev. |
|---|---|
| MEOO | 0.06 ± 0.02 |
| IMIN | 0.05 ± 0.03 |
| IMINH | 0.05 ± 0.03 |
| FORO | 0.046 ± 0.007 |
| FORN | 0.047 ± 0.007 |
| FORC | 0.045 ± 0.007 |
| DMEO | 0.06 ± 0.02 |
| BENC | 0.02 ± 0.02 |
| PRPC | 0.019 ± 0.004 |
| GEHC | 0.03 ± 0.04 |
| MAMN | 0.03 ± 0.01 |
| ACEC | 0.038 ± 0.003 |
| ACEO | 0.03 ± 0.01 |
| TIPO/SWM4O | 0.003 ± 0.003 |

Table S5: SILCS-MC protocols used for each ligand. An * indicates that 250 simulated annealing steps were attempted

| Target | Ligand | FF | Method |
|---|---|---|---|
| IVP | DPQ | C36 | Pose Refinement |
| | | Drude | Pose Refinement |
| GUA | dGUA | C36 | Pose Refinement |
| | | Drude | Pose Refinement* |
| TAR | RBT203 | C36 | Pose Refinement* |
| | | Drude | Pose Refinement |
| | RRM | C36 | Pose Refinement |
| | | Drude | Pose Refinement* |
| | NEOB | C36 | Pose Refinement |
| | | Drude | Pose Refinement |
| HCV | BENZ | C36 | Pose Refinement* |
| | | Drude | Pose Refinement |
| TGQ | MM41 | C36 | Exhaustive Docking |
| | | Drude | Exhaustive Docking |

Table S6: Ligand sampling region dimensions for groove binders in Å.

| | **IVP - DPQ** | | **TAR - rbt203** | | **TAR - RRM** | | **TAR - NeoB** | |
|---|---|---|---|---|---|---|---|---|
| | C36 | Drude | C36 | Drude | C36 | Drude | C36 | Drude |
| SILCS-MC Radius | 10 | 10 | 10 | 10 | 10 | 10 | 10 | 10 |
| Glide (Inner) | 20 × 10 × 15 | 20 × 10 × 15 | 20 × 20 × 20 | 20 × 20 × 20 | 20 × 20 × 20 | 20 × 20 × 20 | 12 × 12 × 12 | 12 × 12 × 12 |
| Glide (Outer) | 30 × 20 × 25 | 30 × 20 × 25 | 40 × 40 × 40 | 37 × 37 × 37 | 40 × 40 × 40 | 37 × 37 × 37 | 27 × 27 × 27 | 27 × 27 × 27 |
| Gnina | 20 × 20 × 20 | 20 × 20 × 20 | 35 × 35 × 35 | 35 × 35 × 35 | 35 × 35 × 35 | 35 × 35 × 35 | 20 × 20 × 20 | 20 × 20 × 20 |

Table S7: Ligand sampling region dimensions for alternative binders in Å.

| | **GUA - dGua** | | **HCV - Benz** | | **TGQ - MM41** | |
|---|---|---|---|---|---|---|
| | C36 | Drude | C36 | Drude | C36 | Drude |
| SILCS-MC Radius | 5 | 5 | 10 | 10 | 10 | 10 |
| Glide (Inner) | 18 × 18 × 18 | 12 × 12 × 12 | 18 × 18 × 15 | 18 × 18 × 15 | 15 × 10 × 8 | 15 × 10 × 8 |
| Glide (Outer) | 38 × 38 × 38 | 32 × 32 × 32 | 38 × 38 × 35 | 38 × 38 × 35 | 35 × 30 × 28 | 35 × 30 × 28 |
| Gnina | 25 × 25 × 25 | 25 × 25 × 25 | 20 × 20 × 20 | 20 × 20 × 20 | 35 × 30 × 28 | 35 × 30 × 28 |

Table S8: Docking and pose refinement results from SILCS-MC: LGFE (kcal/mol), LE (kcal/mol·atom), and RMSD (Å).

| | C36 | | | Drude | | |
|---|---|---|---|---|---|---|
| | LGFE | LE | RMSD | LGFE | LE | RMSD |
| **DPQ** | | | | | | |
| 1 | 1.16 | 0.06 | 3.22 | -3.15 | -0.15 | 3.79 |
| 2 | 1.17 | 0.06 | 2.64 | -3.03 | -0.14 | 3.72 |
| 3 | 1.25 | 0.06 | 3.32 | -2.77 | -0.13 | 3.46 |
| 4 | 1.26 | 0.06 | 3.37 | -2.70 | -0.13 | 2.37 |
| 5 | 1.27 | 0.06 | 3.43 | -2.71 | -0.13 | 2.86 |
| 6 | 1.30 | 0.06 | 3.21 | -2.61 | -0.12 | 2.63 |
| 7 | 1.36 | 0.07 | 2.80 | -2.42 | -0.12 | 3.29 |
| 8 | 1.39 | 0.07 | 2.80 | -2.52 | -0.12 | 2.19 |
| 9 | 1.40 | 0.07 | 2.84 | -2.50 | -0.12 | 2.91 |
| Avg RMSD | 3.38 ± 0.73 | | | 2.99 ± 0.7 | | |
| **RBT203** | | | | | | |
| 1 | -5.96 | -0.24 | 3.49 | -8.03 | -0.32 | 2.50 |
| 2 | -5.94 | -0.24 | 3.31 | -7.79 | -0.31 | 2.82 |
| 3 | -5.80 | -0.23 | 2.96 | -7.75 | -0.31 | 4.32 |
| 4 | -5.27 | -0.21 | 3.95 | -7.52 | -0.30 | 5.65 |
| 5 | -5.17 | -0.21 | 3.75 | -7.52 | -0.30 | 6.07 |
| 6 | -4.72 | -0.19 | 3.36 | -7.26 | -0.29 | 5.22 |
| 7 | -4.70 | -0.19 | 3.54 | -7.14 | -0.29 | 4.41 |
| 8 | -4.59 | -0.18 | 3.87 | -7.03 | -0.28 | 5.62 |
| 9 | -4.50 | -0.18 | 3.12 | -7.03 | -0.28 | 5.83 |
| Avg RMSD | 3.88 ± 1.24 | | | 5.54 ± 1.7 | | |
| **RRM** | | | | | | |
| 1 | 0.28 | 0.00 | 2.11 | -7.14 | -0.04 | 1.49 |
| 2 | 0.29 | 0.00 | 2.00 | -7.12 | -0.04 | 1.33 |
| 3 | 0.46 | 0.01 | 2.24 | -6.87 | -0.04 | 2.14 |
| 4 | 0.79 | 0.01 | 2.35 | -6.84 | -0.04 | 2.15 |
| 5 | 0.87 | 0.01 | 2.22 | -6.80 | -0.04 | 2.70 |
| 6 | 0.87 | 0.01 | 1.76 | -6.80 | -0.04 | 2.77 |
| 7 | 0.91 | 0.01 | 2.52 | -6.69 | -0.04 | 1.75 |
| 8 | 0.96 | 0.01 | 2.52 | -6.69 | -0.04 | 1.27 |
| 9 | 1.14 | 0.01 | 2.03 | -6.69 | -0.04 | 1.37 |
| Avg RMSD | 2.14 ± 0.35 | | | 1.93 ± 0.37 | | |
| **NEOB** | | | | | | |
| 1 | -1.30 | -0.03 | 4.06 | -10.51 | -0.25 | 2.25 |
| 2 | -1.12 | -0.03 | 3.58 | -10.48 | -0.25 | 2.20 |
| 3 | -1.05 | -0.03 | 2.68 | -10.08 | -0.24 | 2.78 |
| 4 | -0.89 | -0.02 | 3.21 | -9.73 | -0.23 | 3.26 |
| 5 | -0.85 | -0.02 | 3.21 | -9.63 | -0.23 | 3.30 |
| 6 | -0.62 | -0.02 | 3.99 | -9.56 | -0.23 | 2.16 |
| 7 | -0.37 | -0.01 | 2.98 | -9.53 | -0.23 | 2.12 |
| 8 | -0.37 | -0.01 | 3.86 | -9.53 | -0.23 | 2.46 |
| 9 | -0.22 | -0.01 | 3.10 | -9.29 | -0.22 | 2.16 |
| Avg RMSD | 3.41 ± 0.48 | | | 2.68 ± 0.42 | | |

| | C36 | | | Drude | | |
|---|---|---|---|---|---|---|
| | LGFE | LE | RMSD | LGFE | LE | RMSD |
| **dGUA** | | | | | | |
| 1 | -7.98 | -0.42 | 1.53 | -11.69 | -0.62 | 1.34 |
| 2 | -7.69 | -0.41 | 2.23 | -11.45 | -0.60 | 1.23 |
| 3 | -7.63 | -0.40 | 1.44 | -11.23 | -0.59 | 1.07 |
| 4 | -7.27 | -0.38 | 2.19 | -11.06 | -0.58 | 1.26 |
| 5 | -7.08 | -0.37 | 1.34 | -10.96 | -0.58 | 1.08 |
| 6 | -7.04 | -0.37 | 1.74 | -10.93 | -0.57 | 1.05 |
| 7 | -6.99 | -0.37 | 1.48 | -10.87 | -0.57 | 0.85 |
| 8 | -6.89 | -0.36 | 1.11 | -10.75 | -0.57 | 1.06 |
| 9 | -6.68 | -0.35 | 1.25 | -10.65 | -0.56 | 1.10 |
| Avg RMSD | 1.59 ± 0.4 | | | 1.14 ± 0.14 | | |
| **BENZ** | | | | | | |
| 1 | -4.72 | -0.20 | 3.49 | -6.39 | -0.27 | 1.89 |
| 2 | -4.70 | -0.20 | 3.51 | -6.33 | -0.27 | 2.15 |
| 3 | -4.61 | -0.19 | 3.57 | -6.06 | -0.25 | 2.20 |
| 4 | -4.51 | -0.19 | 3.35 | -5.83 | -0.25 | 2.52 |
| 5 | -3.45 | -0.14 | 3.60 | -5.68 | -0.24 | 2.51 |
| 6 | -2.98 | -0.12 | 3.74 | -5.46 | -0.23 | 2.67 |
| 7 | -2.54 | -0.11 | 2.90 | -5.46 | -0.23 | 2.27 |
| 8 | -2.50 | -0.10 | 2.94 | -5.45 | -0.23 | 2.11 |
| 9 | -2.39 | -0.10 | 2.94 | -5.46 | -0.23 | 2.67 |
| Avg RMSD | 3.4 ± 0.3 | | | 2.23 ± 0.27 | | |
| **MM41** | | | | | | |
| 1 | -9.33 | -0.16 | 8.75 | -21.10 | -0.35 | 6.85 |
| 2 | -8.98 | -0.16 | 9.41 | -20.96 | -0.35 | 7.97 |
| 3 | -8.64 | -0.14 | 9.98 | -20.67 | -0.35 | 7.59 |
| 4 | -8.60 | -0.14 | 9.58 | -20.57 | -0.34 | 5.49 |
| 5 | -8.47 | -0.14 | 10.00 | -19.97 | -0.33 | 4.24 |
| 6 | -8.46 | -0.14 | 10.10 | -19.57 | -0.32 | 5.02 |
| 7 | -8.27 | -0.14 | 9.84 | -19.44 | -0.32 | 3.83 |
| 8 | -8.17 | -0.14 | 9.88 | -19.44 | -0.32 | 3.86 |
| 9 | -8.15 | -0.14 | 9.66 | -19.35 | -0.32 | 4.56 |
| Avg RMSD | 9.69 ± 0.54 | | | 6.07 ± 1.82 | | |

Table S9: Docking score (kcal/mol), $\Delta G_{\mathrm{MMGBSA}}$ (kcal/mol), and RMSD (Å) from Glide.

| | C36 | | | Drude | | |
|---|---|---|---|---|---|---|
| | Docking Score | $\Delta G_{MMGBSA}$ | RMSD | Docking Score | $\Delta G_{MMGBSA}$ | RMSD |
| **IVP-DPQ** | | | | | | |
| 1 | -6.51 | -32.04 | 4.64 | -6.00 | -35.36 | 4.86 |
| 2 | -6.23 | -35.39 | 4.44 | -5.83 | -36.46 | 4.66 |
| 3 | -6.05 | -32.55 | 4.60 | -5.83 | -32.97 | 4.56 |
| 4 | -5.94 | -34.38 | 4.36 | -5.83 | -29.54 | 5.22 |
| 5 | -5.94 | -34.80 | 4.50 | -5.78 | -33.10 | 5.32 |
| 6 | -5.94 | -34.42 | 4.56 | -5.78 | -33.62 | 5.62 |
| 7 | -5.68 | -31.06 | 4.97 | -4.96 | -31.54 | 5.14 |
| Avg RMSD | 4.69 ± 0.17 | | | 4.82 ± 0.1 | | |
| **TAR-RBT203** | | | | | | |
| 1 | -8.63 | -41.22 | 10.20 | -8.53 | -36.32 | 9.15 |
| 2 | -8.59 | -42.68 | 9.23 | -8.51 | -35.92 | 9.55 |
| 3 | -8.57 | -42.65 | 9.63 | -8.51 | -43.00 | 9.64 |
| 4 | -7.76 | -50.99 | 12.01 | -8.51 | -37.54 | 9.89 |
| 5 | -7.52 | -50.95 | 9.73 | -8.36 | -32.77 | 13.28 |
| 6 | -7.46 | -53.26 | 10.00 | -7.76 | -27.50 | 10.02 |
| 7 | -7.35 | -47.85 | 9.25 | -7.65 | -48.64 | 12.00 |
| 8 | -6.98 | -45.56 | 12.51 | -7.56 | -48.64 | 12.41 |
| Avg RMSD | 10.7 ± 1.74 | | | 10.29 ± 1.6 | | |
| **TAR-RRM** | | | | | | |
| 1 | -8.29 | -68.39 | 12.12 | -10.01 | -52.62 | 13.15 |
| 2 | -7.78 | -37.13 | 13.21 | -7.94 | -46.19 | 12.91 |
| 3 | -7.59 | -57.40 | 9.62 | | | |
| 4 | -6.58 | -58.13 | 9.22 | | | |
| 5 | -6.54 | -58.13 | 8.24 | | | |
| 6 | -6.52 | -57.46 | 8.64 | | | |
| 7 | -6.28 | -57.36 | 8.05 | | | |
| 8 | -5.23 | -57.6 | 8.81 | | | |
| Avg RMSD | 9.92 ± 1.66 | | | 13.03 ± 0.17 | | |
| **TAR-NEOB** | | | | | | |
| 1 | -7.69 | -55.27 | 7.21 | -6.64 | -53.37 | 7.27 |
| 2 | -6.84 | -51.52 | 9.74 | -6.64 | -53.13 | 7.25 |
| 3 | -6.21 | -47.18 | 9.74 | -6.64 | -52.70 | 7.25 |
| 4 | -6.21 | -48.47 | 9.71 | -6.57 | -53.29 | 7.05 |
| 5 | -6.16 | -51.93 | 9.71 | -6.27 | -51.64 | 7.21 |
| 6 | -6.14 | -48.79 | 10.41 | -6.21 | -51.82 | 7.60 |
| 7 | -6.00 | -47.56 | 11.03 | -6.21 | -50.71 | 7.41 |
| 8 | -5.76 | -49.36 | 11.41 | -6.21 | -52.83 | 8.04 |
| 9 | -5.72 | -45.92 | 11.03 | -6.21 | -48.32 | 6.71 |
| Avg RMSD | 9.89 ± 1.96 | | | 7.63 ± 0.48 | | |

| | C36 | | | Drude | | |
|---|---|---|---|---|---|---|
| | Docking Score | $\Delta G_{MMGBSA}$ | RMSD | Docking Score | $\Delta G_{MMGBSA}$ | RMSD |
| **GUA-dGua** | | | | | | |
| 1 | -6.25 | -24.08 | 10.74 | -7.64 | -33.97 | 1.06 |
| 2 | -6.24 | -38.04 | 13.94 | -7.60 | -35.95 | 1.66 |
| 3 | -6.14 | -40.21 | 14.58 | -7.59 | -34.02 | 1.06 |
| 4 | -6.11 | -41.81 | 14.58 | -7.57 | -33.92 | 1.07 |
| 5 | -6.08 | -41.61 | 15.76 | -7.55 | -33.82 | 1.08 |
| 6 | -6.06 | -42.38 | 14.66 | -7.55 | -33.92 | 1.08 |
| 7 | -5.78 | -22.07 | 11.82 | -7.35 | -34.42 | 1.27 |
| 8 | -5.65 | -62.02 | 13.64 | -7.29 | -34.42 | 1.07 |
| 9 | -5.65 | -62.02 | 13.92 | -6.04 | -27.41 | 7.47 |
| Avg RMSD | 13.26 ± 1.82 | | | 4.63 ± 3.31 | | |
| **HCV-BENZ** | | | | | | |
| 1 | -9.84 | -38.09 | 6.08 | -10.48 | -55.95 | 1.74 |
| 2 | -9.49 | -34.04 | 5.21 | -10.43 | -58.72 | 1.65 |
| 3 | -9.41 | -30.59 | 5.89 | -10.27 | -50.70 | 1.74 |
| 4 | -9.40 | -38.67 | 5.22 | -10.23 | -53.02 | 1.64 |
| 5 | -9.29 | -47.62 | 5.95 | -9.82 | -55.37 | 1.70 |
| 6 | -9.25 | -47.16 | 6.08 | -9.79 | -55.97 | 1.74 |
| 7 | -7.29 | -40.55 | 13.97 | -9.52 | -58.12 | 2.23 |
| 8 | -7.25 | -41.62 | 13.95 | | | |
| 9 | -7.09 | -57.16 | 13.93 | | | |
| Avg RMSD | 9.62 ± 4.6 | | | 2.23 ± 0.38 | | |
| **TGQ-MM41** | | | | | | |
| 1 | -9.20 | -46.58 | 8.58 | -9.40 | -72.11 | 9.10 |
| 2 | -9.42 | -50.20 | 8.72 | -8.98 | -71.10 | 7.63 |
| 3 | -9.87 | -55.78 | 9.05 | -8.97 | -70.45 | 9.30 |
| 4 | -8.97 | -55.78 | 9.56 | -8.95 | -69.60 | 6.85 |
| 5 | -8.97 | -55.78 | 9.56 | -8.95 | -69.57 | 6.73 |
| 6 | -8.97 | -55.78 | 9.56 | -8.95 | -67.45 | 6.63 |
| 7 | -8.97 | -55.78 | 9.62 | -8.87 | -65.79 | 6.30 |
| 8 | -8.95 | -55.78 | 9.46 | -8.86 | -65.29 | 6.03 |
| 9 | -8.42 | -51.84 | 9.70 | -8.17 | -65.32 | 6.83 |
| Avg RMSD | 9.46 ± 1 | | | 8.79 ± 1.36 | | |

Table S10: Binding affinity (kcal/mol), intramolecular energy (kcal/mol), and RMSD (Å) for all poses from Gnina.

| | C36 | | | Drude | | |
|---|---|---|---|---|---|---|
| | Binding Affinity | Intramolecular Energy | RMSD | Binding Affinity | Intramolecular Energy | RMSD |
| **IVP-DPQ** | | | | | | |
| 1 | -5.57 | -0.36 | 4.61 | -6.22 | -0.33 | 7.39 |
| 2 | -6.02 | -0.31 | 6.82 | -5.67 | -0.37 | 4.52 |
| 3 | -5.12 | -0.37 | 5.49 | -5.80 | -0.33 | 4.69 |
| 4 | -5.48 | -0.33 | 8.33 | -5.34 | -0.33 | 7.29 |
| 5 | -5.79 | -0.31 | 5.59 | -5.75 | -0.38 | 6.94 |
| 6 | -5.40 | -0.35 | 6.65 | -5.06 | -0.31 | 6.61 |
| 7 | -5.49 | -0.32 | 7.58 | -5.46 | -0.31 | 6.50 |
| 8 | -6.13 | -0.32 | 5.55 | -5.28 | -0.36 | 5.77 |
| 9 | -5.85 | -0.36 | 9.28 | -5.13 | -0.35 | 6.09 |
| Avg RMSD | | | **6.66 ± 1.52** | | | **6.20 ± 1.04** |
| **TAR-RBT203** | | | | | | |
| 1 | -6.00 | -0.03 | 14.16 | -5.62 | -0.66 | 13.48 |
| 2 | -5.69 | -0.95 | 12.89 | -5.88 | -0.88 | 12.08 |
| 3 | -4.67 | -0.84 | 16.54 | -5.75 | -1.36 | 13.16 |
| 4 | -5.74 | -0.70 | 8.85 | -5.16 | -0.65 | 11.90 |
| 5 | -5.67 | -1.43 | 10.53 | -5.56 | -1.06 | 14.90 |
| 6 | -5.68 | -0.90 | 14.89 | -5.01 | -0.85 | 10.81 |
| 7 | -6.03 | -0.79 | 12.03 | -5.21 | -1.05 | 10.44 |
| 8 | -5.90 | -1.09 | 10.75 | -5.73 | -0.66 | 15.03 |
| 9 | -5.42 | -1.13 | 14.17 | -5.30 | -0.62 | 4.63 |
| Avg RMSD | | | **12.76 ± 2.44** | | | **11.83 ± 3.14** |
| **TAR-RRM** | | | | | | |
| 1 | -7.59 | -7.43 | 8.29 | -8.09 | -8.53 | 15.83 |
| 2 | -7.82 | -11.37 | 9.21 | -7.14 | -11.25 | 8.14 |
| 3 | -7.37 | -7.53 | 9.83 | -7.24 | -9.96 | 7.88 |
| 4 | -7.64 | -3.44 | 12.50 | -7.28 | -9.54 | 12.96 |
| 5 | -6.74 | -8.13 | 8.20 | -7.74 | -9.78 | 16.05 |
| 6 | -6.98 | -8.00 | 8.65 | -6.78 | -9.39 | 17.00 |
| 7 | -6.86 | -9.37 | 10.07 | -6.82 | -11.25 | 7.71 |
| 8 | -7.10 | -11.71 | 12.99 | -7.82 | -9.62 | 15.71 |
| 9 | -7.62 | -10.82 | 13.21 | -7.50 | -10.97 | 8.11 |
| Avg RMSD | | | **10.33 ± 2.04** | | | **12.15 ± 4.12** |
| **TAR-NEOB** | | | | | | |
| 1 | -8.76 | -1.51 | 3.67 | -8.15 | -2.04 | 6.32 |
| 2 | -7.00 | -1.35 | 9.69 | -7.47 | -2.15 | 9.39 |
| 3 | -7.83 | -1.39 | 5.51 | -7.10 | -2.49 | 10.49 |
| 4 | -7.96 | -3.05 | 9.45 | -7.06 | -0.99 | 6.08 |
| 5 | -7.18 | -1.57 | 9.43 | -8.39 | -1.89 | 3.72 |
| 6 | -7.90 | -2.67 | 8.95 | -7.76 | -0.50 | 5.23 |
| 7 | -8.91 | -1.57 | 9.33 | -7.86 | -0.66 | 9.26 |
| 8 | -7.19 | -3.08 | 4.11 | -7.79 | -2.43 | 9.54 |
| 9 | -8.90 | -0.48 | 5.94 | -6.95 | -2.31 | 6.50 |
| Avg RMSD | | | **7.34 ± 2.50** | | | **7.39 ± 2.33** |

| | C36 | | | Drude | | |
|---|---|---|---|---|---|---|
| | Binding Affinity | Intramolecular Energy | RMSD | Binding Affinity | Intramolecular Energy | RMSD |
| **GUA-dGua** | | | | | | |
| 1 | -10.13 | -0.52 | 7.09 | -7.86 | -0.18 | 7.75 |
| 2 | -8.97 | 0.16 | 8.11 | -7.07 | -0.55 | 11.32 |
| 3 | -9.02 | 0.02 | 7.32 | -7.45 | -0.30 | 12.29 |
| 4 | -8.47 | -0.46 | 3.19 | -6.93 | -0.42 | 11.49 |
| 5 | -8.42 | -0.45 | 6.19 | -7.21 | 0.28 | 10.83 |
| 6 | -8.58 | -0.50 | 11.66 | -6.30 | -0.34 | 11.29 |
| 7 | -7.95 | -0.40 | 9.35 | -7.21 | -0.48 | 10.45 |
| 8 | -8.63 | -0.49 | 8.59 | -6.90 | -0.54 | 11.84 |
| 9 | -7.52 | -0.49 | 12.39 | -6.63 | -0.11 | 10.53 |
| Avg RMSD | | | **8.21 ± 2.79** | | | **10.86 ± 1.31** |
| **HCV-BENZ** | | | | | | |
| 1 | -8.65 | 2.13 | 1.88 | -8.68 | 0.33 | 1.15 |
| 2 | -8.35 | 0.43 | 1.94 | -8.69 | 0.22 | 1.76 |
| 3 | -8.17 | 2.30 | 1.36 | -6.91 | 0.19 | 2.15 |
| 4 | -8.39 | -0.52 | 0.79 | -7.22 | -0.63 | 1.34 |
| 5 | -8.06 | -0.54 | 1.99 | -6.64 | 1.56 | 5.53 |
| 6 | -7.65 | 0.47 | 5.29 | -6.90 | 0.46 | 5.19 |
| 7 | -7.97 | 0.33 | 5.62 | -6.97 | -0.02 | 5.59 |
| 8 | -7.04 | -0.67 | 2.25 | -7.66 | -0.32 | 5.56 |
| 9 | -7.83 | 0.15 | 5.62 | -7.36 | -0.60 | 2.79 |
| Avg RMSD | | | **2.97 ± 1.95** | | | **3.45 ± 1.97** |
| **TGQ-MM41** | | | | | | |
| 1 | -6.36 | 0.34 | 12.88 | -6.59 | -2.44 | 9.33 |
| 2 | -6.45 | -2.38 | 12.84 | -6.06 | -2.22 | **6.12** |
| 3 | -6.51 | -2.71 | 12.69 | -6.01 | -2.44 | 6.76 |
| 4 | -7.61 | -1.98 | 11.98 | -6.60 | -1.98 | 12.16 |
| 5 | -7.04 | -1.42 | **11.76** | -6.96 | -2.34 | 10.38 |
| 6 | -7.21 | -2.22 | 11.91 | -6.05 | -2.65 | 9.69 |
| 7 | -6.84 | -3.03 | 11.79 | -6.43 | -1.92 | 10.42 |
| 8 | -6.76 | -1.90 | 12.40 | -6.14 | -2.20 | 11.68 |
| 9 | -6.45 | -2.16 | 12.28 | -6.00 | -2.29 | 10.27 |
| Avg RMSD | | | **12.28 ± 0.45** | | | **9.64 ± 2.03** |

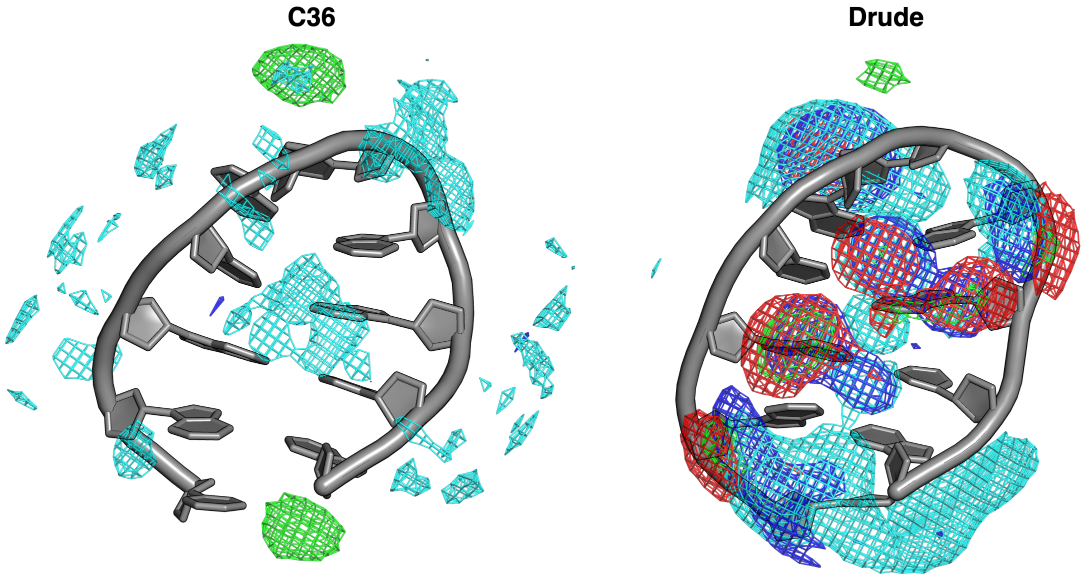


Figure S2: SILCS FragMaps for DDX. FragMaps are shown as apolar (light green), GFE level -1.0; HB Donor (blue), GFE level -1.0; HB Acc (red) , GFE level -1.0; AceO (orange), GFE level -1.0; MeoO (tan), GFE level -1.0; MamN (cyan), GFE level -1.2. Bottom view is shown as it is the primary binding site for a known ligand.

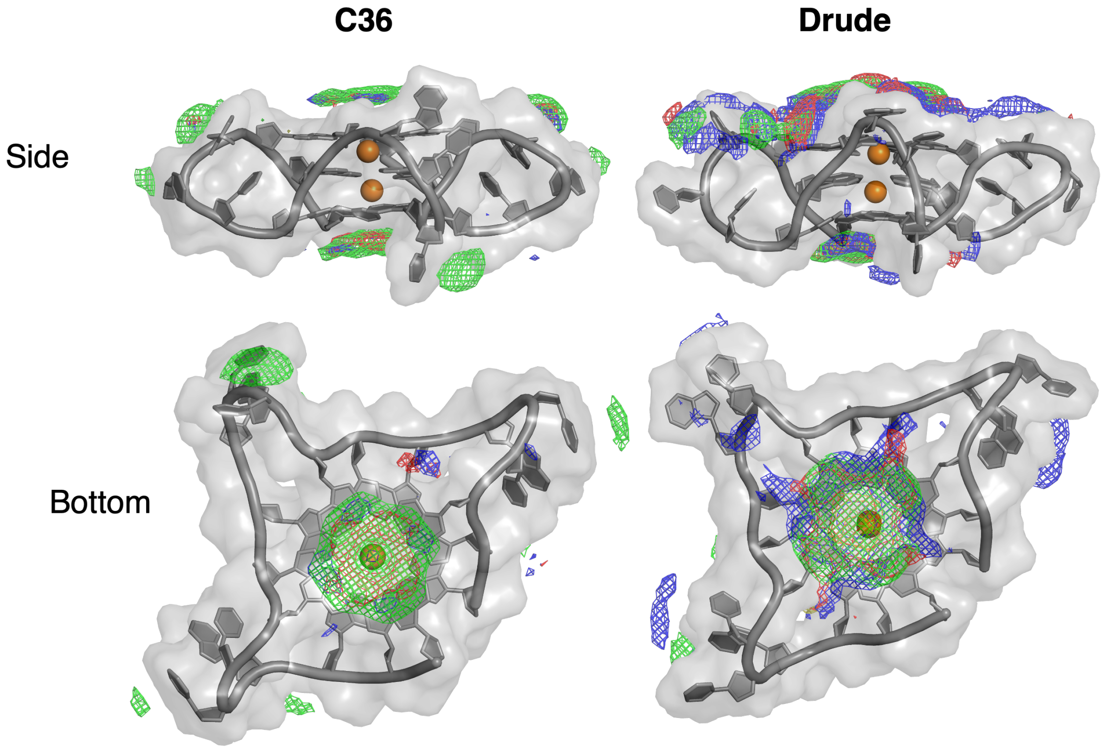


Figure S3: SILCS FragMaps for TGQ. FragMaps are shown as apolar (light green), GFE level -0.8; HB Donor (blue), GFE level -0.8; HB Acc (red) , GFE level -0.8; AceO (orange), GFE level -0.8; and MeoO (tan), GFE level -0.8. Bottom view is shown as it is the primary binding site for a known ligand.

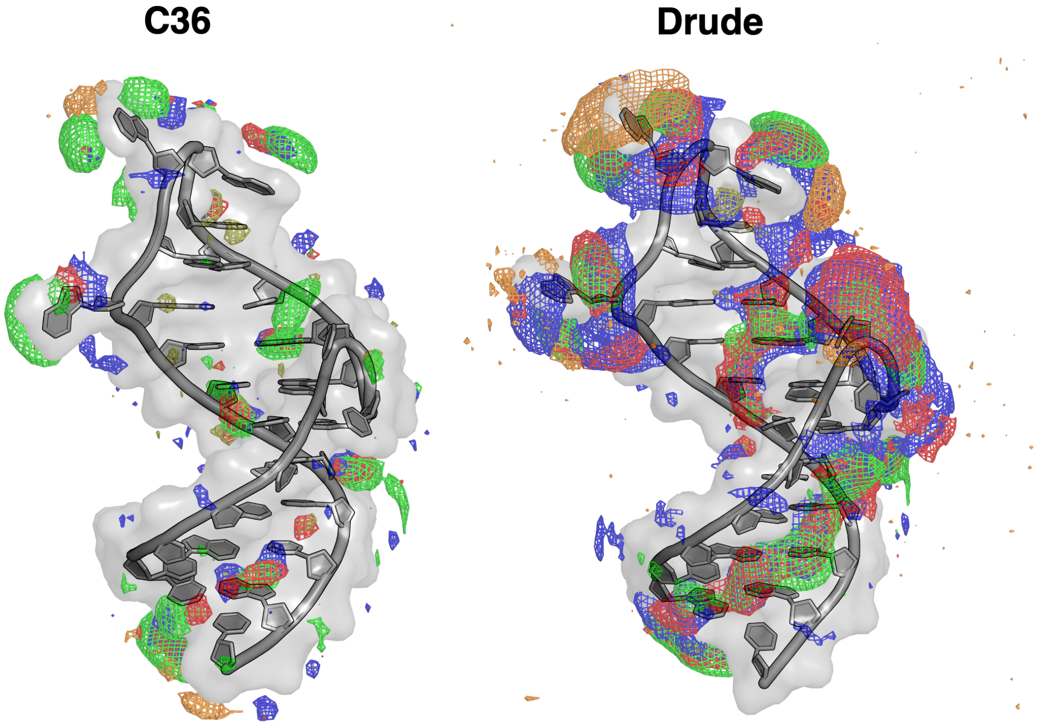


Figure S4: SILCS FragMaps for TAR. FragMaps are shown as apolar (light green), GFE level -0.5; HB Donor (blue), GFE level -0.5; HB Acc (red) , GFE level -0.5; AceO (orange), GFE level -0.5; and MeoO (tan), GFE level -0.5.

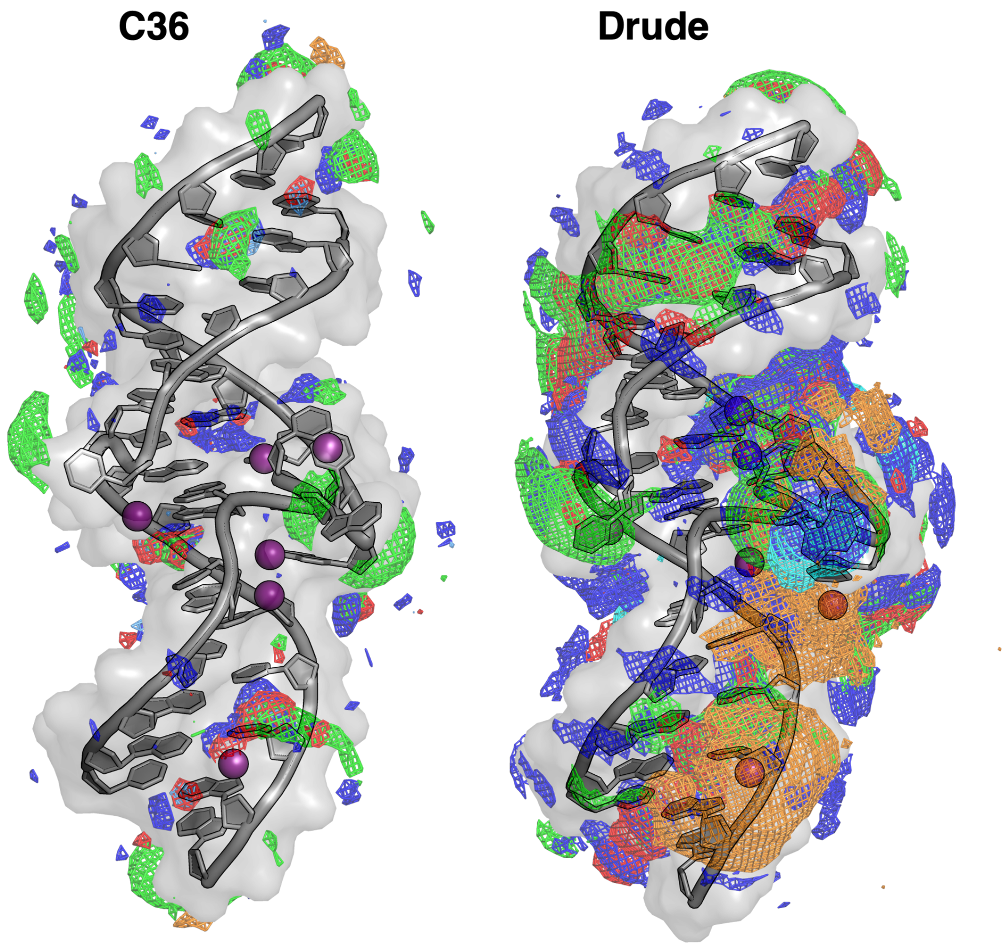


Figure S5: SILCS FragMaps for HCV. FragMaps are shown as apolar (light green), GFE level -0.5; HB Donor (blue), GFE level -0.5; HB Acc (red) , GFE level -0.5; AceO (orange), GFE level -0.5; and MeoO (tan), GFE level -0.5.

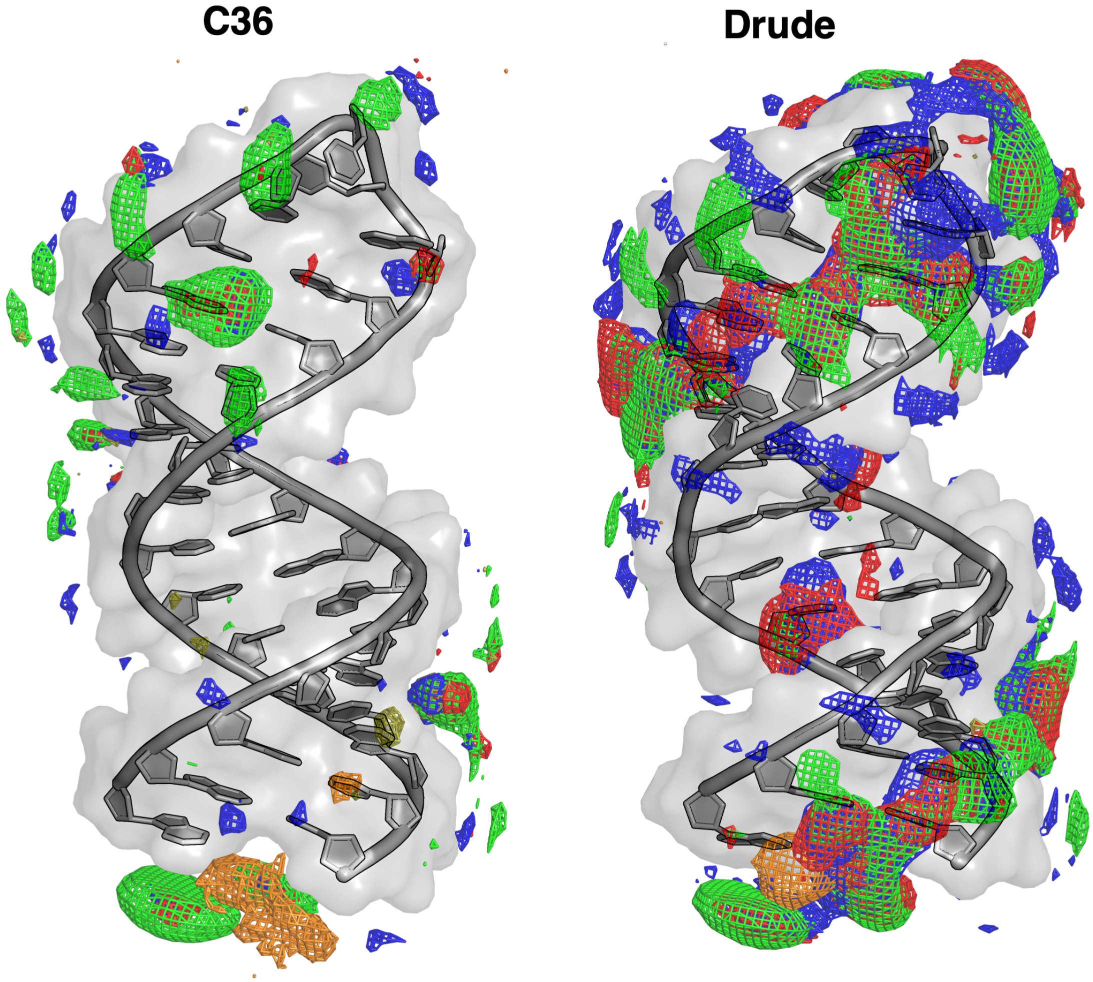


Figure S6: SILCS FragMaps for IVP. FragMaps are shown as apolar (light green), GFE level -0.5; HB Donor (blue), GFE level -0.5; HB Acc (red) , GFE level -0.5; AceO (orange), GFE level -0.5; and MeoO (tan), GFE level -0.5.

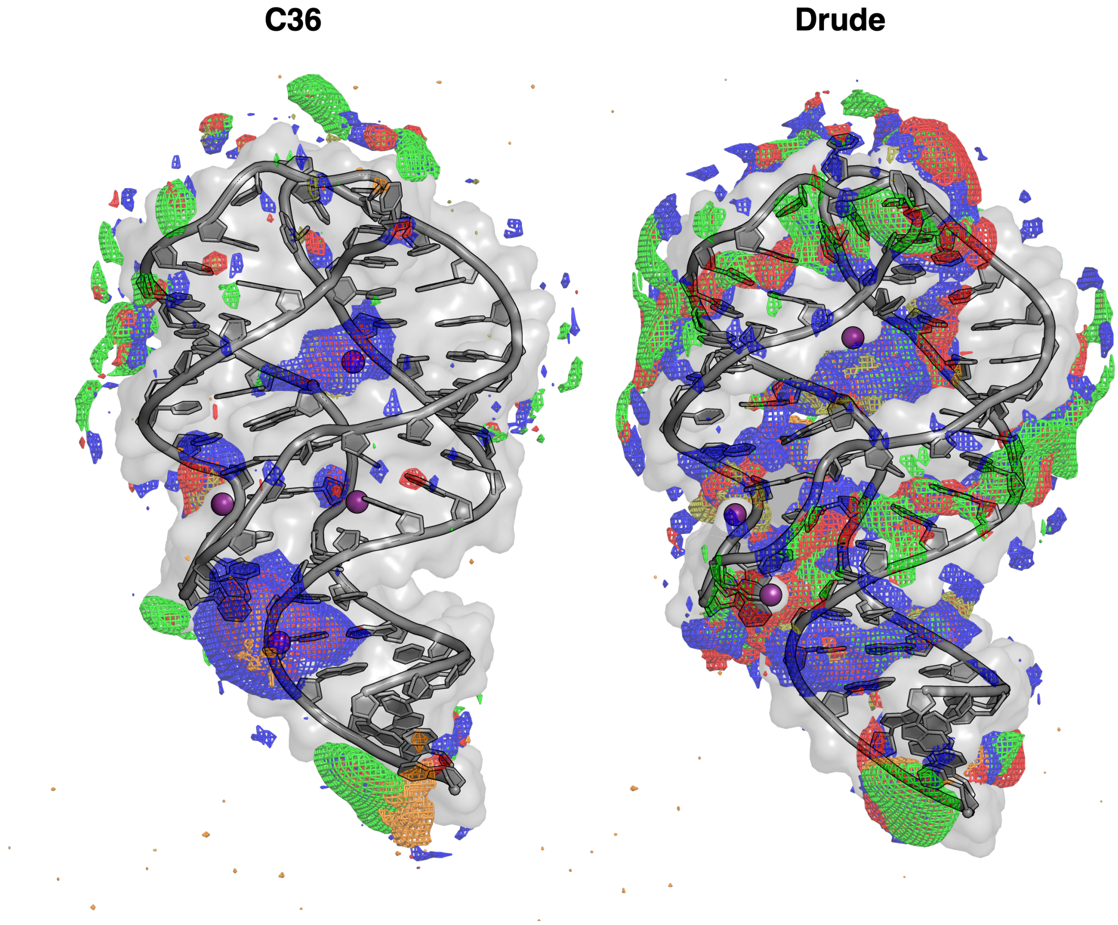


Figure S7: SILCS FragMaps for GUA. FragMaps are shown as apolar (light green), GFE level -0.5; HB Donor (blue), GFE level -0.5; HB Acc (red) , GFE level -0.5; AceO (orange), GFE level -0.5; and MeoO (tan), GFE level -0.5.

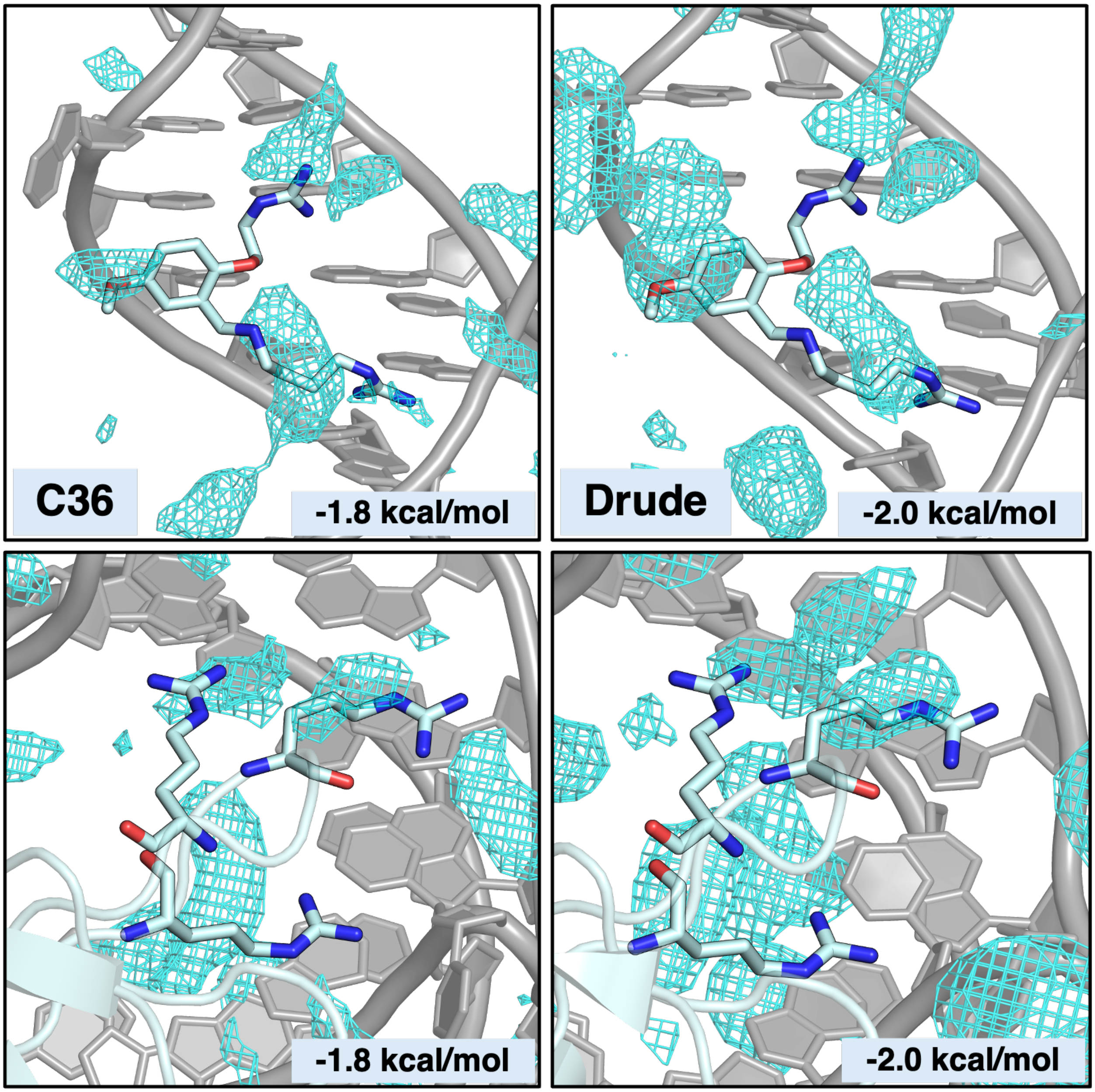


Figure S8: RBT203 (Top) and RRM (Bottom) within their known binding sites on TAR. Positively charged maps are are localized near the guanidinium interaction sites for both ligands at high interaction favorability. Methylammonium maps are shown at the labeled GFE.

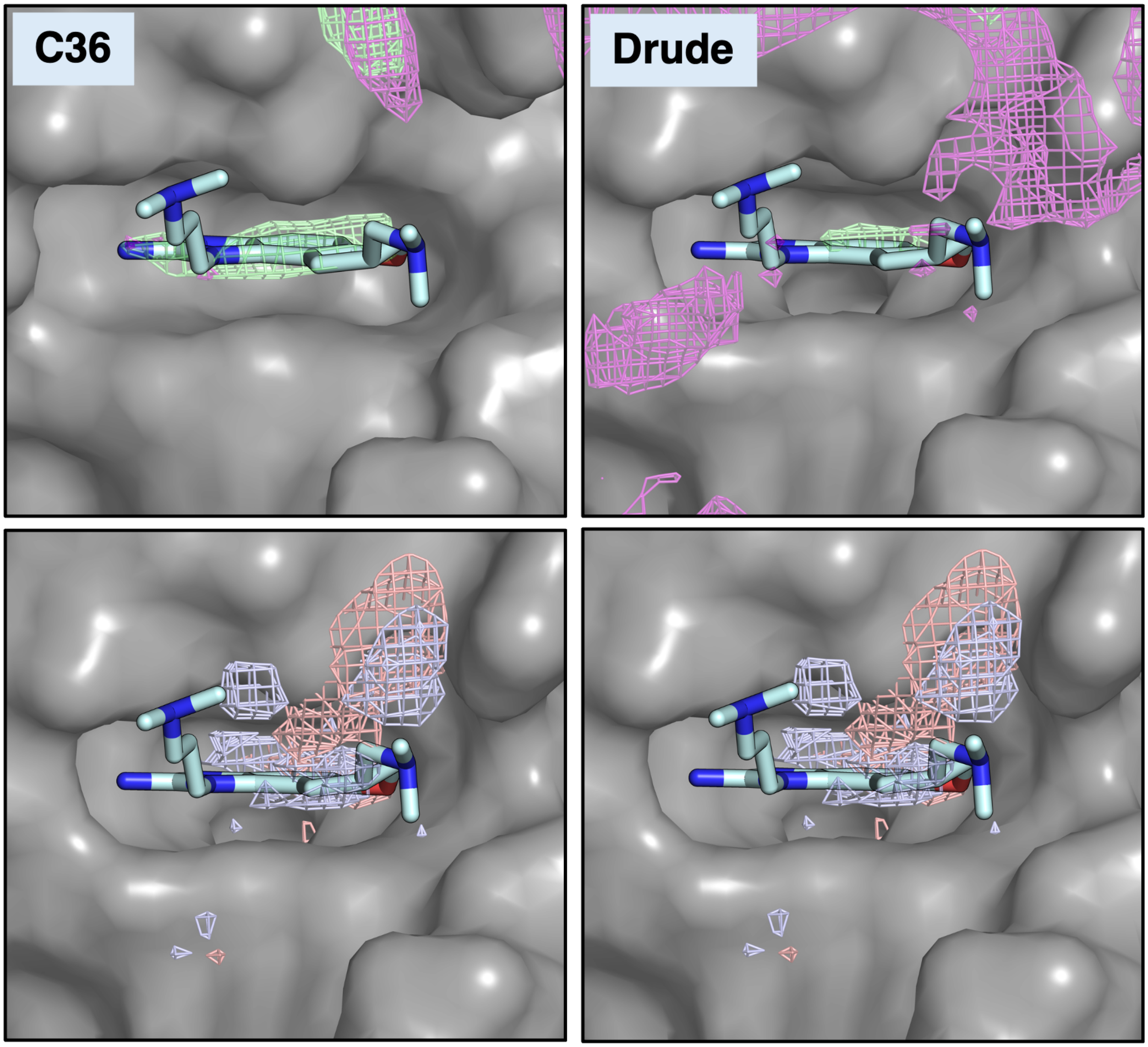


Figure S9: Comparison of C36 and Drude FragMaps for benzene, propane, and imidazole within the intercalation site of HCV. FragMaps are overlaid on the crystal pose and are shown as BENX C (purple), GFE level -0.2; PRPX C (lime green), GFE level -0.5; IMIN N (light blue) , GFE level -1.4; IMIN NH (salmon), GFE level -1.4. The BENX maps indicate no sampling within the binding pocket for both C36 and Drude.

C36 Drude

**A. DPQ**

| RMSD (Å) | | |
|---|---|---|
| Method | C36 | Drude |
| ◻ SILCS | 2.64 | 2.19 |
| ■ Glide | 4.36 | 4.69 |
| ■ Gnina | 4.61 | 4.52 |

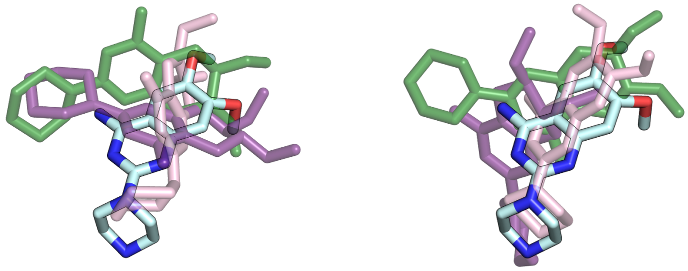

**B. RBT203**

| RMSD (Å) | | |
|---|---|---|
| Method | C36 | Drude |
| ◻ SILCS | 3.04 | 2.50 |
| ■ Glide | 6.76 | 9.16 |
| ■ Gnina | 8.85 | 4.63 |

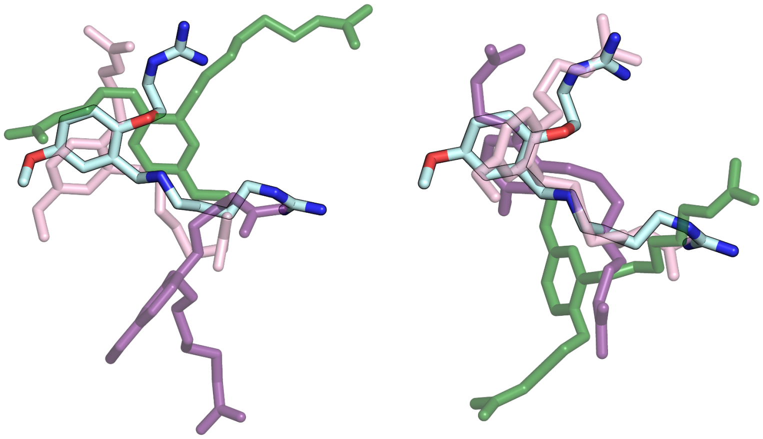

**C. NEOB**

| RMSD (Å) | | |
|---|---|---|
| Method | C36 | Drude |
| ◻ SILCS | 2.68 | 2.10 |
| ■ Glide | 5.13 | 6.02 |
| ■ Gnina | 3.67 | 5.23 |

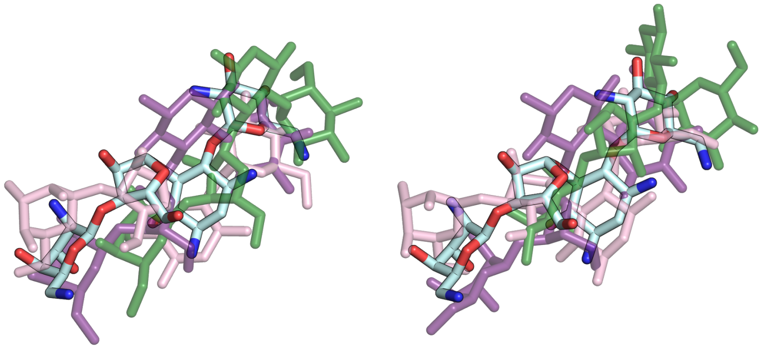

Figure S10: Lowest RMSD poses for groove binding ligands A) DPQ, B) RBT203, and C) NEOB. SILCS-MC (light pink), Glide (green), and Gnina (purple) are overlaid on the original experimental pose (cyan).

C36 Drude

**A. dGUA**

| RMSD (Å) | | |
|---|---|---|
| Method | C36 | Drude |
| ◻ SILCS | 1.11 | 0.85 |
| ◼ Glide | 4.36 | 4.69 |
| ◼ Gnina | 3.19 | 7.75 |

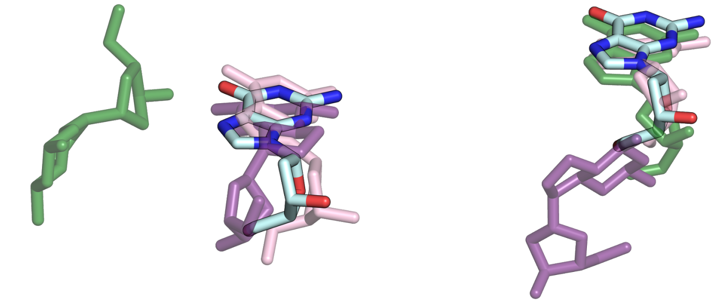

**B. BENZ**

| RMSD (Å) | | |
|---|---|---|
| Method | C36 | Drude |
| ◻ SILCS | 2.90 | 1.89 |
| ◼ Glide | 1.58 | 1.74 |
| ◼ Gnina | 0.79 | 1.15 |

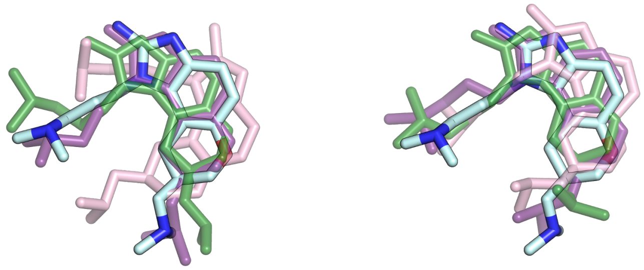

**C. MM41**

| RMSD (Å) | | |
|---|---|---|
| Method | C36 | Drude |
| ◻ SILCS | 8.75 | 4.56 |
| ◼ Glide | 8.51 | 6.35 |
| ◼ Gnina | 11.76 | 6.12 |

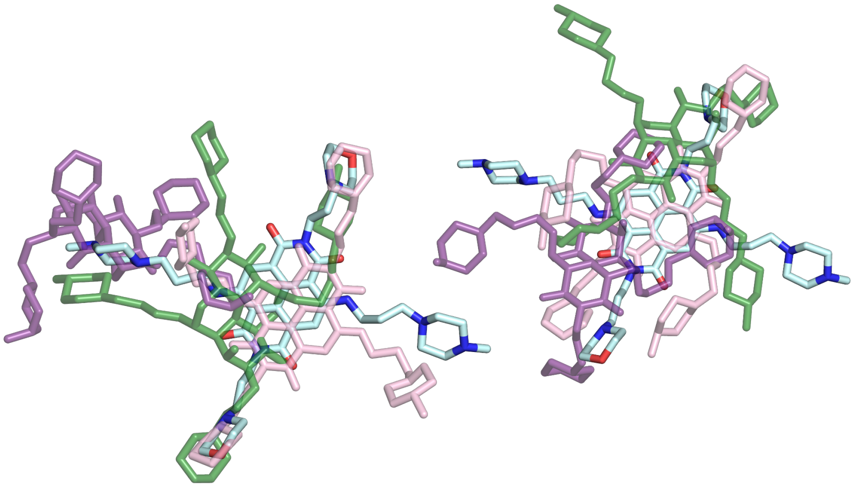

Figure S11: Lowest RMSD poses for alternative binding ligands A) dGUA, B) BENZ, and C) MM41. SILCS-MC (light pink), Glide (green), and Gnina (purple) are overlaid on the crystal pose (cyan).

(3) Hwang, W. et al. CHARMM at 45: Enhancements in Accessibility, Functionality, and Speed. *The Journal of Physical Chemistry B 128*, 9976–10042.

(4) *Kirk-Othmer Encyclopedia of Chemical Technology*, 4th ed.; John Wiley and Sons: New York, NY, 1994; Vol. 11; p 960, Series published 1991–Present.

(5) Chickos, J. S.; Acree, W. E. Thermochemical Properties of Formamide Revisited: New Experiment and Quantum Mechanical Calculations. *Journal of Chemical & Engineering Data* **2011**, *56*, 4183–4187.

(6) Leader, G. R. The Dielectric Constant of Formamide1. *Journal of the American Chemical Society* **1951**, *73*, 856–857, _eprint: https://doi.org/10.1021/ja01146a513.

(7) Budavari, S., Ed. *The Merck Index: An Encyclopedia of Chemicals, Drugs, and Biologicals*, 12th ed.; Merck and Co., Inc.: Whitehouse Station, NJ, 1996; p 718.

(8) Goel, H.; Yu, W.; Ustach, V. D.; Aytenfisu, A. H.; Sun, D.; MacKerell, A. D. Impact of electronic polarizability on protein-functional group interactions. *Physical Chemistry Chemical Physics 22*, 6848–6860.
